# CD300e is a driver of the immunosuppressive tumor microenvironment and colorectal cancer progression via macrophage reprogramming

**DOI:** 10.1101/2024.09.01.610700

**Authors:** Annica Barizza, Stefania Vassallo, Laura Masatti, Mattia Laffranchi, Sofia Giacometti, Silvia Lonardi, Mattia Bugatti, Sara Coletta, Simone Pizzini, William Vermi, Matteo Fassan, Gaya Spolverato, Silvano Sozzani, Enrica Calura, Gaia Codolo

## Abstract

Colorectal cancer (CRC) progression is shaped by the tumor microenvironment, particularly tumor-associated macrophages (TAMs), which often adopt immunosuppressive functions. CD300e, a myeloid receptor involved in immune regulation, has an uncharacterized role in CRC. Here, we show that CD300e is selectively upregulated in tumor-infiltrating monocytes and macrophages, driving a suppressive phenotype marked by impaired antigen presentation. In vitro cocultures of patient-derived tumor organoids and human monocytes revealed that tumor-derived signals induce CD300e expression and promote a protumorigenic macrophage profile. Using CD300e knockout mice in AOM/DSS and MC38 CRC models, we found that CD300e loss reduced tumor burden, enhanced MHC expression on TAMs, and improved T-cell responses. Transcriptomic and functional analyses demonstrated that CD300e-deficient macrophages exhibit increased phagocytosis, upregulated antigen presentation, and greater support for T-cell proliferation and cytotoxicity. Adoptive transfer confirmed that macrophage-intrinsic CD300e expression is sufficient to suppress T-cell function and promote tumor growth. Our findings identify CD300e as a critical regulator of macrophage-mediated immune suppression in CRC and a potential target for reprogramming TAMs to enhance immunotherapy.

## Introduction

Colorectal cancer (CRC) is the most common malignancy of the gastrointestinal tract and the second most common cause of death related to cancer[1,2]. Emerging evidence has revealed the pivotal role of signals derived from the tumor microenvironment (TME) in the development and progression of CRC. The TME comprises not only malignant cells but also infiltrating immune cells, stromal cells, vasculature, and a milieu of secreted factors and extracellular matrix components [3,4]. Within this complex ecosystem, immune cells, particularly T lymphocytes and myeloid cells, play a dual role, either promoting tumor elimination or enabling immune escape and tumor growth, depending on their functional orientation.

Recent studies have revealed a strong correlation between the immune landscape of the TME and clinical outcomes in CRC patients [5,6]. The development of the Immunoscore system, which quantifies cytotoxic T-cell infiltration in tumor tissue, has underscored the prognostic value of immune composition in CRC, often surpassing conventional TNM staging in predicting patient survival [7,8]. Despite this, many CRC patients exhibit poor responses to immune checkpoint blockade therapies, suggesting the presence of dominant immunosuppressive mechanisms within the TME that inhibit effective antitumor immunity.

Among the immune populations that support cancer growth in the TME, tumor-associated macrophages (TAMs) are the most abundant [9–11]. Although macrophages have the potential to mount a robust antitumor response, the phenotype of TAMs becomes progressively skewed toward an immunosuppressive, tumor-promoting profile as cancer progresses [12,13]. This reprogramming is driven by tumor-derived signals and results in a macrophage population that suppresses cytotoxic lymphocyte function, promotes angiogenesis, remodels the extracellular matrix, and facilitates metastatic dissemination [14]. For example, TAMs can inhibit the recruitment of CD8^+^ T cells to the tumor site, impair antigen presentation, engage immune checkpoints pathways, and foster the expansion of regulatory T (Treg) cells [15,16]. High TAM infiltration in CRC has been consistently associated with worse clinical outcomes, poor responses to immunotherapy, and tumor resistance to conventional treatments [17–20].

Given their central role in modulating the immune contexture of tumors, TAMs have emerged as promising therapeutic targets. However, clinical strategies aimed at depleting or reprogramming TAMs have yielded limited efficacy, emphasizing the need to better understand the molecular signals that govern TAM behaviour in the TME [21–23].

One emerging candidate is CD300e, an immune receptor expressed predominantly on myeloid cells, including monocytes and macrophages, in both humans and mice [24,25]. Initially classified as an activating receptor due to its association with pro-inflammatory signaling [26], CD300e has more recently been implicated in negative regulation of immune responses. Notably, CD300e engagement impairs macrophage-mediated T-cell activation by inhibiting antigen presentation, thus promoting immune evasion [27,28]. Moreover, upregulation of CD300e has been linked to a regenerative or immunosuppressive macrophage phenotype in various pathological contexts [29–31]. Despite its expression in several tumor types, including CRC, the functional role of CD300e in tumor immunity remains uncharacterized. Intriguingly, elevated CD300e expression correlates with poor prognosis across multiple cancer types, positioning it as a potential immune regulatory checkpoint [32,33].

In this study, we identify CD300e as a previously unrecognized driver of macrophage-mediated immune suppression in CRC. Through analysis of patient-derived samples, we show that CD300e is selectively upregulated in TAMs with a protumoral, immunosuppressive phenotype. Using patient-derived organoid–monocyte co-cultures, we demonstrate that tumor-derived cues are sufficient to induce CD300e expression and polarize macrophages toward a tumor-promoting state. In murine CRC models, genetic ablation of CD300e reduced tumor burden, enhanced MHC expression on TAMs, and restored T-cell-mediated antitumor responses. Our findings reveal a novel immunosuppressive pathway mediated by CD300e and suggest that therapeutic targeting of this receptor may reprogram TAMs and enhance the efficacy of immunotherapy in CRC.

## Materials and methods

### Subjects

Surgical resections of tumor (CRC) and normal (NC) colonic tissue were collected from 8 CRC patients who underwent curative-intent surgery (General Surgery 3, University Hospital of Padua) following ethical standards, the Declaration of Helsinki, and national and international guidelines. Written informed consent was obtained from the individual patients, and the Ethical Committee of Padua University Hospital approved the protocol (Protocol Number: P448). The inclusion criteria for the enrolled patients were as follows: patients with histologically confirmed primary adenocarcinoma of the colon, patients aged >18 years, and patients who provided written informed consent. Patients with a known history of hereditary colorectal cancer syndrome and patients who had undergone neoadjuvant treatments were excluded. The patients’ characteristics are detailed in Table 1.

**Table 1.**
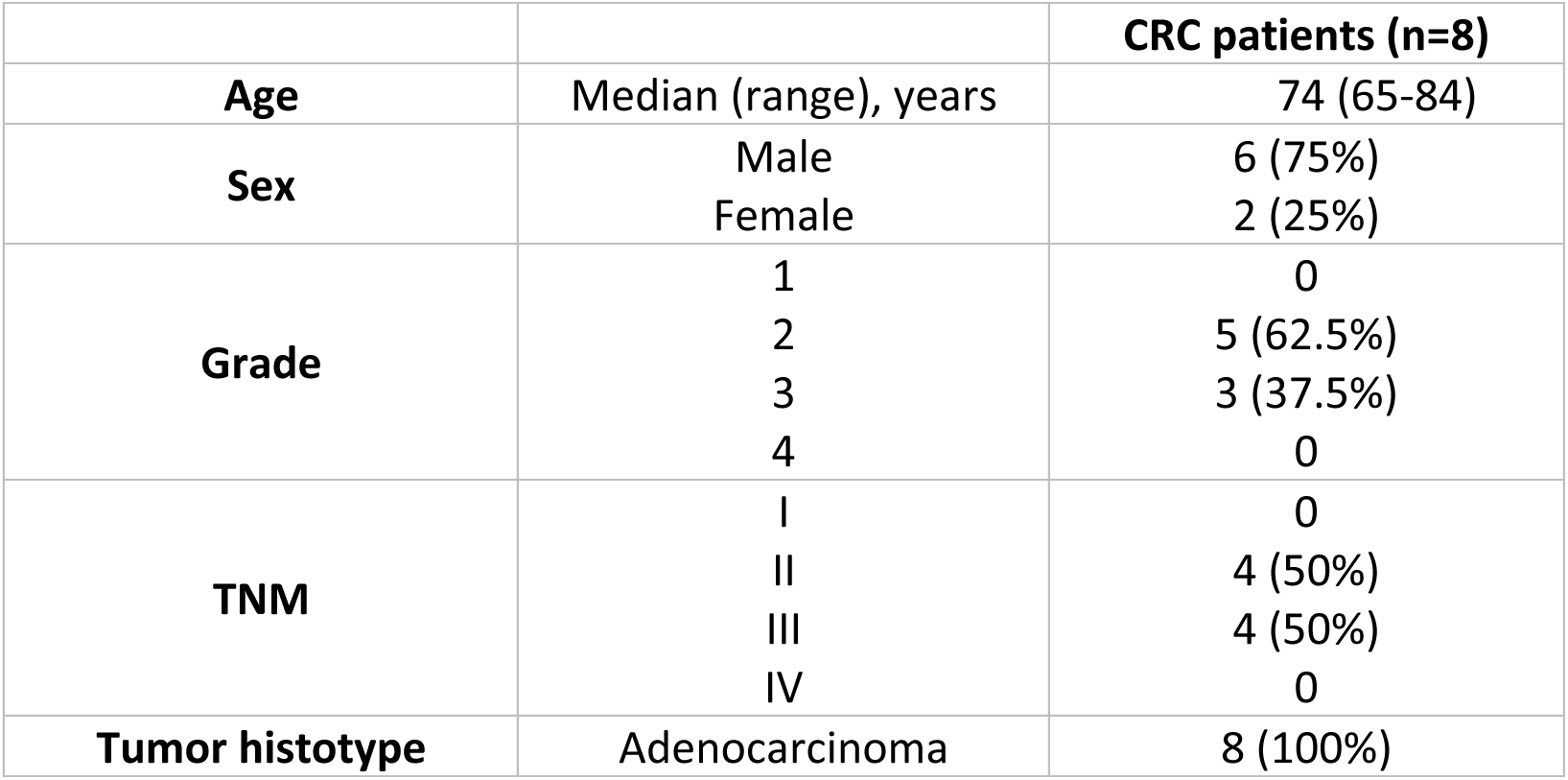
Clinicopathological characteristics of CRC patients. CRC: colorectal cancer; TNM: pathological tumor, node, metastasis stage.

The peripheral blood mononuclear cells utilized in this study were derived from buffy coats obtained from healthy blood donors and anonymously provided by the Transfusion Centre of the University Hospital of Padua because of voluntary and informed blood donation for transfusions. Written informed consent for the use of the buffy coats for research purposes was obtained from blood donors by the Transfusion Centre.

Further experimental details are available in the online supplemental materials and methods section.

### Tumor and normal colon organoids

#### Generation of patient-derived organoids

Patient-derived tumor colon organoids (TCOs) were generated from surgical resections of CRC patients’ primary tumor tissue, as previously reported by Sato et al. [34]. The tumor tissue was minced into small pieces (2–4 mm) and digested in DMEM supplemented with 2% FBS, 2 mg/mL collagenase II (Sigma Aldrich, St. Louis, MO, USA), 100 μg/mL DNAse I (Roche, Basel, Switzerland), 100 U/mL penicillin, and 100 µg/mL streptomycin for 1 h at 37°C under constant rolling. The cell suspension was filtered through a 100 μm filter, washed in Advanced DMEM/F12 (Gibco, Thermo Fisher Scientific, Waltham, MA, USA) supplemented with 2 mM GlutaMAX (Gibco), 1 mM HEPES (Corning, Corning, NY, USA), 100 U/mL penicillin, and 100 µg/mL streptomycin, and seeded in Growth Factor Reduced (GFR) Matrigel^TM^ (Corning) (2000 cells/μL).

Patient-derived normal colon organoids (NCOs) were generated from colonic crypts obtained from CRC patient-matched normal mucosa. Mucosal tissue was fragmented into small pieces (2–4 mm) and incubated in 3 mM EDTA or 0.5 mM dithiothreitol (DTT) in PBS with 2% FBS for 40 minutes at 4°C under constant rolling to allow for crypt release. The crypt suspension was filtered through a 100 μm filter, washed with Advanced DMEM/F12 supplemented with 2 mM GlutaMAX, 1 mM HEPES, 100 U/mL penicillin, and 100 µg/mL streptomycin, and seeded in GFR Matrigel^TM^.

#### Organoid culture

Organoids were cultured in tumor or normal colon organoid medium (advanced DMEM/F12 supplemented with 2 mM GlutaMAX, 10 mM HEPES, 100 U/mL penicillin, 100 µg/mL streptomycin, 20% R-Spondin1 conditioned medium (RCM), 10% Noggin conditioned medium (NCM), 10 mM nicotinamide (Sigma Aldrich), 1X B27 supplement (Gibco), 1 mM N-acetyl-L-cysteine (Cayman Chemical, Ann Arbor, MI, USA), 50 ng/mL EGF (Corning), 10 μM Y-27632 (ROCK inhibitor, Selleckchem, Houston, TX, USA), 500 nM A83-01 (Tocris Bioscience, Bristol, UK), 3 μM SB202190 (Sigma Aldrich), and 100 μg/mL primocin (InvivoGen, San Diego, CA, USA)). Fifty percent Wnt-3a-conditioned medium (WCM) was added to normal colon organoid medium. Gastrin I (10 nM, Sigma Aldrich) and prostaglandin E2 (10 nM, Sigma Aldrich) were added to the medium of both patient-derived TCO and NCO. WCM, RCM and NCM were obtained from Wnt-3a-, R-spondin1-, and Noggin-producing cell lines, respectively. Organoids were passaged 6–7 days after seeding and then once a week.

### Monocyte purification, macrophage differentiation and polarization

Human monocytes were purified from buffy coats of healthy donors as reported elsewhere [35] or via a Pan Monocyte Isolation Kit (Miltenyi Biotec, Bergisch Gladbach, North Rhine-Westphalia, Germany) following the manufacturer’s instructions.

Mouse bone marrow cells were isolated to purify monocytes and differentiate into macrophages in vitro. Bone marrow cells were obtained by flushing femurs and tibias with RPMI medium supplemented with 10% FBS, 100 U/mL penicillin, and 100 µg/mL streptomycin. The cells were filtered through a 70 μm filter, and red blood cells were lysed via BD PharmLyse Lysing Buffer (Becton Dickinson, Franklin Lakes, NJ, USA). Monocytes were purified from bone marrow cells via a Monocyte Isolation Kit (BM), mouse (Miltenyi Biotec).

To differentiate macrophages, bone marrow cells were seeded in Petri dishes in RPMI medium supplemented with 10% FBS, 100 U/mL penicillin, 100 µg/mL streptomycin, and 10% conditioned medium obtained from the L929 cell line, which contained macrophage–colony stimulating factor (M-CSF). Macrophage differentiation was achieved 7 days after seeding.

### Cell lines

MC38 mouse adenocarcinoma cells were cultured in DMEM supplemented with 10% FBS, 100 U/mL penicillin, and 100 µg/mL streptomycin. When ∼80% confluence was reached, the cells were injected into the mice. L929 mouse fibroblasts were cultured in DMEM supplemented with 10% FBS, 100 U/mL penicillin, and 100 µg/mL streptomycin. L929 conditioned medium was used to differentiate bone marrow-derived macrophages in vitro. The Wnt-3a-, R-spondin1- and Noggin-producing cell lines were cultured in DMEM supplemented with 10% FBS, 100 U/mL penicillin, and 100 µg/mL streptomycin, according to the datasheet.

### Cocultures of monocytes/macrophages and organoids

Human monocytes purified from buffy coats were cocultured together with either patient-derived TCOs or NCOs. Three to four days after passage, the organoid culture medium was removed, and 2.5×10^5^ monocytes resuspended in DMEM supplemented with 10% FBS, 4 mM HEPES and 50 μg/mL gentamycin were added to each organoid well. After five days of coculture, monocytes/macrophages were harvested and analyzed by flow cytometry and qRT‒PCR; coculture supernatants were collected for analysis of cytokine content.

### Western blot

Western blotting was performed as reported elsewhere [36]. Briefly, human NC and CRC tissues from surgical resections of CRC patients were lysed in RIPA buffer (50 mM Tris-HCl, pH 7.4; 150 mM NaCl; NP-40, 1%; Na-deoxycholate, 0.5%; SDS, 0.1%; 2 mM EDTA; 50 mM NaF; 1 mM Na3VO4; 1 mM EGTA; and 2% PMSF) with a protease inhibitor cocktail (Millipore, Burlington, MA, USA) via TissueLyser II (Qiagen, Hilden, North Rhine-Westphalia, Germany) to homogenize the tissue. Proteins were quantified with a BCA protein assay kit (Thermo Fisher Scientific) according to the manufacturer’s instructions. Thirty micrograms of total protein from each sample was separated by SDS‒PAGE and transferred to PVDF membranes (Millipore). The antigens were revealed via a rabbit anti-vinculin antibody (1:5000, clone 42H89L44, Invitrogen, Waltham, MA, USA) and a rabbit anti-human CD300e polyclonal antibody (1:1000, Sigma Aldrich), followed by an HRP-conjugated anti-rabbit IgG secondary antibody (Millipore). The blots were developed with enhanced chemiluminescence substrate (EuroClone, Pero, Italy), and protein bands were detected via an ImageQuant LAS 4000 (GE Healthcare, Chicago, IL, USA).

### Mouse experimental models

#### Animals

The mice were housed at the Department of Biology, University of Padua Animal Facility. All protocols were approved by the ethical committee of the Italian Ministry of Health (Protocols number 978/2020-PR, 386/2022-PR and 858/2023-PR). The mice were housed in a temperature (22–24°C) and humidity-controlled (50%) colony room and maintained on a 12 h light/dark cycle with standard food and freely available water and environmental enrichment.

CD300e total KO and CD300e conditional KO mouse models were generated on a C57BL/6J background at the Transgenic Core Facility of the University of Copenhagen via the CRISPR/Cas9 system in mouse zygotes. Heterozygous male mice carrying a deletion encompassing exon 2 of the *Cd300E* gene in all body tissues were used as founders of the new CD300e^-/-^ mouse line. Similarly, heterozygous male mice with LoxP site insertions flanking exon 2 were used as founders for generating the new CD300e^fl/fl^ mouse line, which was crossed with LysM Cre mice to generate a conditional KO mouse lacking CD300e expression on myeloid cells (CD300e^LysM^). Wild-type (CD300e^+/+^) C57Bl/6j mice were used as controls for CD300e^-/-^ mice, and CD300e^fl/fl^ mice were used as controls for CD300e^LysM^. All experiments were conducted on age- and sex-matched 8-- to 10--week-old mice housed in the same animal facility.

#### AOM/DSS-induced colorectal cancer

To induce the development of colitis-associated colorectal cancer, male mice were injected intraperitoneally with a single dose of 10 mg/kg azoxymethane (AOM, Sigma Aldrich). Three days later, the mice received 2% dextran sodium sulfate (DSS, molecular weight: 36–50 kDa, MP Biomedicals, Irvine, CA, USA) in their drinking water for 5 days, followed by 16 days of recovery in regular water. This cycle of DSS treatment was repeated two more times, and the mice were sacrificed 4 weeks after the last DSS administration. The clinical course of colitis was evaluated by monitoring the body weight of the mice during the experiment and the presence of diarrhea and blood after the DSS cycles.

#### Subcutaneous tumor model and macrophage adoptive transfer

To establish a syngeneic tumor model, a 100 µL tumor cell suspension containing 10^6^ MC38 cells in a mixture of phosphate-buffered saline (PBS) and GFR Matrigel^TM^ (1:1 ratio) was subcutaneously (s.c.) engrafted into the right flank of male mice.

For macrophage adoptive transfer, macrophages isolated from CD300e^+/+^ and CD300e^-/-^ mice, as well as primary bone marrow-derived macrophages (BMDMs) isolated from CD300e^+/+^ and CD300e^-/-^ mice, were polarized toward a TAM profile with ng/ml mIL-4 20 and 20 ng/ml mIL-13 (Miltenyi Biotec) and mixed with MC38 cells at a ratio of 1:2, and 7.5×10^5^ total cells were s.c. injected into CD300e^+/+^ recipient mice or CD300e^-/-^, as previously reported [37].

The tumor dimensions were measured once the tumors were palpable. Tumor growth was monitored three times per week with a digital caliper, and the tumor volume was calculated via the formula 0.5 × L × W^2^, with L and W being the length and width, respectively. The mice were euthanized on the 21^st^ day after MC38 inoculation or when the tumors reached 1.5 cm in diameter. Once excised, the final volume of the tumor was accurately calculated by multiplying the length, width, and height (L × W × H).

### FITC-dextran permeability assays

The mice were starved overnight and then orally administered 600 mg/kg FITC-dextran (4 kDa, Sigma‒Aldrich) through gavage. After 3 h, blood was collected in EDTA-treated tubes from the submandibular vein and centrifuged at 3000 rpm for 15 min to obtain plasma. A mixture of plasma and PBS (1:10 ratio) in a total volume of 100 µL was transferred to a flat-bottom black 96-well plate, and the fluorescence intensity of FITC-dextran (excitation: 483 nm; emission: 530 nm) was measured via an Infinite 200Pro plate reader (Tecan, Männedorf, Switzerland). The plasma concentration of FITC-dextran was calculated via a standard curve obtained from serial dilutions of stock FITC-dextran (100 µg/ml) in mouse plasma.

### Flow cytometry

Flow cytometry samples were acquired via an LSRFortessa X-20 Cell Analyzer (BD Biosciences, Franklin Lakes, NJ, USA), and the data were analyzed via FlowJo version 10.3 (Tree Star, Ashland, OR, USA).

#### Human monocyte-derived macrophages

After 5 d of coculture with NCO or TCO, human monocyte-derived macrophages were harvested from culture wells with 5 mM Na-EDTA in PBS and incubated for 10 min at room temperature with a human Fc block (BD Biosciences). The cells were stained with human monocyte/macrophage panel antibodies as detailed in Table 2.

**Table 2:**
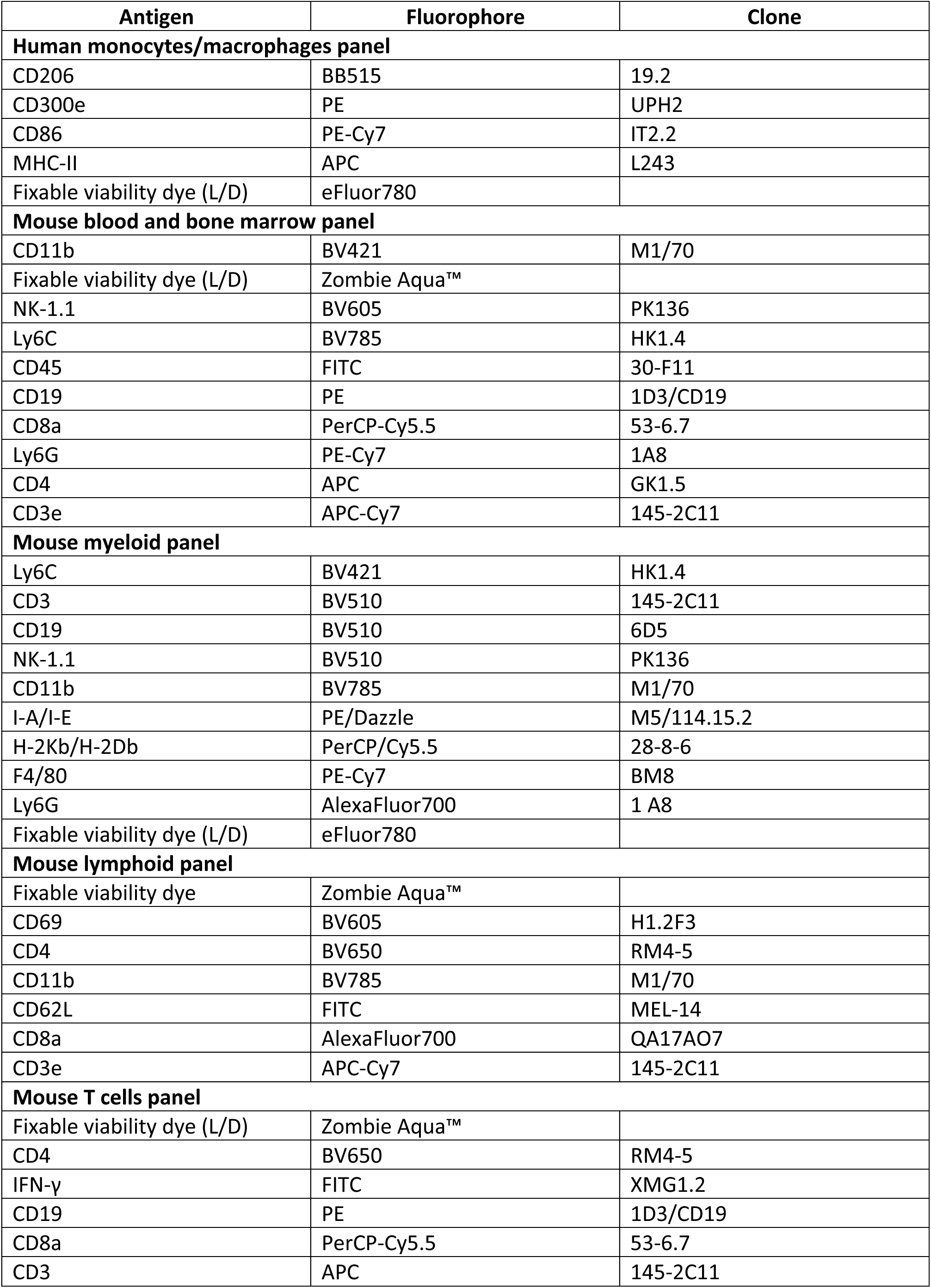
Detailed list of antibody panels used for flow cytometry analysis.

| Antigen | Fluorophore | Clone |
| --- | --- | --- |
| **Human monocytes/macrophages panel** |  |  |
| CD206 | BB515 | 19.2 |
| CD300e | PE | UPH2 |
| CD86 | PE-Cy7 | IT2.2 |
| MHC-II | APC | L243 |
| Fixable viability dye (L/D) | eFluor780 |  |
| **Mouse blood and bone marrow panel** |  |  |
| CD11b | BV421 | M1/70 |
| Fixable viability dye (L/D) | Zombie Aqua™ |  |
| NK-1.1 | BV605 | PK136 |
| Ly6C | BV785 | HK1.4 |
| CD45 | FITC | 30-F11 |
| CD19 | PE | 1D3/CD19 |
| CD8a | PerCP-Cy5.5 | 53-6.7 |
| Ly6G | PE-Cy7 | 1A8 |
| CD4 | APC | GK1.5 |
| CD3e | APC-Cy7 | 145-2C11 |
| **Mouse myeloid panel** |  |  |
| Ly6C | BV421 | HK1.4 |
| CD3 | BV510 | 145-2C11 |
| CD19 | BV510 | 6D5 |
| NK-1.1 | BV510 | PK136 |
| CD11b | BV785 | M1/70 |
| I-A/I-E | PE/Dazzle | M5/114.15.2 |
| H-2Kb/H-2Db | PerCP/Cy5.5 | 28-8-6 |
| F4/80 | PE-Cy7 | BM8 |
| Ly6G | AlexaFluor700 | 1 A8 |
| Fixable viability dye (L/D) | eFluor780 |  |
| **Mouse lymphoid panel** |  |  |
| Fixable viability dye | Zombie Aqua™ |  |
| CD69 | BV605 | H1.2F3 |
| CD4 | BV650 | RM4-5 |
| CD11b | BV785 | M1/70 |
| CD62L | FITC | MEL-14 |
| CD8a | AlexaFluor700 | QA17A07 |
| CD3e | APC-Cy7 | 145-2C11 |
| **Mouse T cells panel** |  |  |
| Fixable viability dye (L/D) | Zombie Aqua™ |  |
| CD4 | BV650 | RM4-5 |
| IFN-γ | FITC | XMG1.2 |
| CD19 | PE | 1D3/CD19 |
| CD8a | PerCP-Cy5.5 | 53-6.7 |
| CD3 | APC | 145-2C11 |

#### Murine blood and bone marrow cells

Peripheral blood from 8-week-old mice was collected in EDTA-treated tubes and incubated with BD PharmLyse Lysing Buffer (BD Biosciences) to lyse erythrocytes. Blood cells or bone marrow cells, which were isolated as previously described, were incubated for 10 minutes at 4°C with TruStain FcX™ PLUS (clone S17011E, BioLegend, San Diego, CA, USA) to saturate the Fc receptors. The cells were stained with mouse blood/bone marrow panel antibodies as detailed in Table 2.

#### Colonic adenomas and subcutaneous tumors

Colonic adenomas and subcutaneous tumors were isolated and cut into small pieces of 2–4 mm in Petri dishes. The minced tissues were transferred into GentleMACS Tubes (Miltenyi Biotec) containing DMEM with 1 mg/mL collagenase IV (Sigma‒Aldrich) and 0.5 mg/mL DNase I. GentleMACS Tubes were attached to a GentleMACS Octo Dissociator (Miltenyi Biotec), and the tumors were maintained for 40 minutes at 37°C under continuous rotation. The resulting cell suspension was filtered through a 70 µm strainer and incubated with TruStain FcX™ PLUS. To characterize the tumor-infiltrating myeloid lineage, the cells were stained with mouse myeloid panel antibodies as detailed in Table 2. To characterize the tumor-infiltrating lymphoid lineage, the cells were stained with mouse lymphoid panel antibodies as detailed in Table 2.

#### T cells isolated from subcutaneous tumors and lymph nodes

Tumor-infiltrating lymphocytes were purified from dissociated MC38-derived tumors via the EasySep Mouse T-Cell Isolation Kit (STEMCELL Technologies, Vancouver, Canada) according to the manufacturer’s instructions. Local lymph nodes of MC38 tumor-bearing mice and mesenteric lymph nodes of AOM/DSS-treated mice were collected, placed on a 70 µm filter and mechanically disrupted via a syringe plunger.

A total of 0.5×10^6^ tumor-infiltrating lymphocytes, as well as T cells isolated from local and mesenteric lymph nodes, were seeded in 48-well plates precoated with purified anti-mouse CD3 (1:50 in PBS; clone 145-2c11; BioLegend) in RPMI supplemented with 10% FBS, 100 U/mL penicillin, and 100 µg/mL streptomycin containing Ultra-LEAF-purified anti-mouse CD28 antibody (1:1000, clone 37.51, BioLegend). After overnight stimulation, BD GolgiStop protein transport inhibitor (containing monensin) (1:10 in RPMI, 10% FBS, 100 U/mL penicillin, 100 µg/mL streptomycin; BD Biosciences) was added, and the mixture was incubated for 5 h. The cells were subsequently incubated with TruStain FcX™ PLUS, and the cells were then stained with mouse T-cell panel antibodies as detailed in Table 2.

For intracellular IFN-γ detection, the cells were fixed and permeabilized with a Cytofix/Cytoperm™ Fixation/Permeabilization Kit (BD Biosciences); FMO tubes were prepared as negative controls.

### Immunosuppressive activity assay

The immunosuppressive activity of F4/80^+^ tumor-infiltrating cells was evaluated by determining the proliferation of allogenic CellTrace™-labeled splenic T cells. Briefly, T cells were stained with 0.5 μM CellTrace™ Violet Cell Proliferation Kit (Invitrogen, Molecular Probes) according to the manufacturer’s instructions, activated by coating with 1 μg/mL anti-CD3 and 5 μg/mL soluble anti-CD28 (BioLegend) mAb and cocultured in flat-bottom 96-well plates at a 1:3 ratio with F4/80^+^ macrophages isolated from CD300e^+/+^ and CD300e^-/-^ tumors with anti-F4/80 MicroBeads UltraPure following the manufacturer’s instructions (Miltenyi Biotec). After four days at 37°C and 5% CO2, T-cell proliferation was assessed by staining the cells with an anti-CD3 APC mAb (BioLegend) for flow cytometry analysis and analyzing them with a BD™ LSRII flow cytometer (BD Biosciences). The T-cell proliferation index was analyzed with FlowJo 10.3 and calculated as the total number of divisions divided by the number of cells that went into division.

### Histology

#### Hematoxylin and eosin

Hematoxylin and eosin (H&E) staining of liver, spleen, and colon samples was performed on formalin-fixed, paraffin-embedded (FFPE) tissue sections. Slices were hydrated, stained with Mayer’s hematoxylin (Bio-Optica, Milan, Italy) and Eosin Y 1% aqueous solution (Bio-Optica) and dehydrated in ethanol. Images were acquired with a Leica DM6 microscope (Leica Biosystems, Nussloch, Germany) via LasX software.

#### Immunohistochemistry

Immunohistochemistry of matched NC and CRC tissues from CRC patients was performed in collaboration with the University of Brescia. FFPE tissue sections were sequentially immunostained as described previously [38]. Briefly, MHC-II (clone V1030, dilution 1:500; Biomeda, Foster City, CA, USA) was revealed via Novolink Polymer (Leica Biosystems). The slides were then decoloured with ethanol, and the previous antibody was stripped with a β-mercaptoethanol/SDS solution in a water bath preheated at 56°C. Double immunohistochemistry using anti-CD163 (dilution 1:80, clone 10D6, Thermo Fisher Scientific) and polyclonal rabbit anti-CD300e (Sigma Aldrich) was performed on the same sections. The first was revealed using Mach4AP Polymer (Biocare Medical, Concord, CA, USA) and Ferangi Blue as a chromogen, and the second was revealed using Novolink Polymer (Leica Biosystems) followed by diaminobenzidine (DB). The two digital slides were synchronized via ImageScope, and pictures of the same area were taken to highlight the same cells sequentially stained for MHC-II/CD300e and CD163. The cells were quantified via optical counting via the ImageScope count tool.

### ELISA

The cytokine content in the coculture supernatants was evaluated with ELISA kits specific for human IL-6 (Thermo Fisher Scientific) and IL-10 (Thermo Fisher Scientific) following the manufacturer’s instructions and an Infinite F200 Pro plate reader (Tecan).

### RNA extraction and qRT‒PCR

Total RNA was extracted with TRIzol reagent (Thermo Fisher Scientific) according to the manufacturer’s protocol. When working with human or murine colon tissues, the tissue was first disrupted via TissueLyser II (Qiagen). The RNA was quantified via a NanoDrop 1000 spectrophotometer (NanoDrop, Wilmington, DE, USA). One microgram of total RNA was reverse transcribed via a high-capacity cDNA reverse transcription kit (Applied Biosystems, Waltham, MA, USA) following the manufacturer’s instructions. Five nanograms of cDNA was used for qRT‒PCR (15 ng when working with tissues), which was performed with PowerUp SYBR Green master mix (Thermo Fisher Scientific). qRT‒PCR was performed in a QuantStudio5 Real-Time PCR System (Thermo Fisher Scientific) according to the following cycle: 95°C for 5 min, 95°C for 15 s, and 60°C for 1 min for 40 cycles. The experiments were performed with at least three technical replicates. For each sample, the data were normalized to the endogenous reference genes 18S and/or β-actin. The sequences of the primers used are listed in Supplementary Table 1.

### Bulk RNA sequencing, single-cell RNA sequencing and transcriptomic analysis

#### RNA sequencing of monocytes

For the bulk RNA sequencing (RNA-seq) experiment, RNA extracted from BM monocytes was analyzed with a 2100 Bioanalyzer (Agilent Technologies, Santa Clara, CA, USA) to assess RNA quality and integrity. Samples were used only if the RNA integrity number (RIN) was greater than 7. Library construction and sequencing were performed by the NGS Facility at the Department of Biology, University of Padua. RNA-seq libraries were constructed via a Stranded mRNA Prep Kit (Illumina, San Diego, CA, USA), and sequencing was performed on a NovaSeq 6000 instrument (Illumina) via a NovaSeq 6000 S4 Reagent Kit v1.5 300 cycles (Illumina). The reads from the RNA-seq data were mapped and quantified with RSEM software (v1.3.1, genome sequence GRCm39, annotations from Ensembl version 109.149, [39]). Transcripts with fewer than 10 counts in at least 60% of the samples per class were filtered out. Differentially expressed genes (DEGs) were computed via edgeR (version 3.42.4, false discovery rate (FDR)<0.05; [39,40]) to compare CD300e^+/+^ and CD300e^-/-^ cells. Gene Ontology pathway enrichment analysis of DEGs was performed with the R package clusterProfiler (version 4.8.3; [41,42]). Pathways with an FDR < 0.05 were considered significant and included in the discussion. Data are available in Supplementary Table 2.

#### Public single-cell RNA sequencing dataset of human colorectal cancer

The human single-cell RNA-seq (scRNA-seq) dataset was downloaded from the GEO (GSE178341) and was provided by Pelka et al. [43]. Only the cells labeled as myeloid by the original authors were included in the downstream analysis. Low-quality cells (gene count <=200 or >=4000) and cells with >=20% mitochondrial gene expression were removed. The filtered raw count matrix was processed via the Seurat package (v5.3.0, [44]); normalizeData and ScaleData functions were applied with standard options. Highly variable genes (HVGs, n=2000) were obtained via the FindVariableFeatures function and informed principal component analysis (PCA). Each patient cell was integrated via the Harmony R package (v0.1, [45]). Harmony integration was performed with batchID as a variable, and the resulting reduction was used for graph-based clustering (resolution 0.5) and uniform manifold approximation and projection (UMAP). The marker genes per cluster were obtained via the FindAllMarkers function (only.pos = TRUE, min.pct = 0.25, min.diff.pct = 0.2. Data are available in Supplementary Table 3.

Pseudotime analysis utilized Monocle3 (v1.3.1, [46]) on only the identified monocyte and macrophage populations. The starting point was set to the Mono_a cluster. All the analyses were conducted in R (version 4.4.3, 2025-02-28 ucrt Trophy Case) via RStudio.

#### Single-cell RNA sequencing of CD300e^+/+^ and CD300e^-/-^ mouse adenomas

CD45^+^ cells from three CD300e^+/+^ and three CD300e^-/-^ mice were isolated and independently labeled via single-cell labeling with a BD single-cell multiplexing kit (BD Biosciences, #633793); next, the six samples were pooled prior to FACS sorting. CD11b^+^ myeloid cells were FACS-sorted into L/D^-^CD3e^-^ CD19-CD11b^+^ cells and processed via single-cell capture and cDNA synthesis with the BD Rhapsody Express single-cell analysis system following the manufacturer’s protocol (BD Biosciences). The resulting cDNA libraries were prepared via mRNA whole-transcriptome analysis (WTA) and a sample tag library preparation protocol (BD Biosciences). Libraries were sequenced on a NovaSeq 6000 sequencer (Illumina).

The raw data were aligned via the BD Rhapsody Sequence Analysis CWL pipeline v2.2.1, with RhapRef_Mouse_WTA_2023-02 used as a reference genome. The h5mu to Seurat pipeline v2.2.1 was subsequently used for conversion (https://openpipelines.bio/). Data processing and normalization were performed through the package Seurat (v5.3.0). The single-cell expression matrix was filtered (700 < nFeature_RNA < 5500 percent.mt < 25%, and 1000 < nCount_RNA < 100000). PCA was computed via the 2000 highly variable features obtained via the FindVariableFeatures function. Graph-based unsupervised clustering and UMAP were performed by selecting the first top 10 PCs.

The marker genes for each cluster were identified via the FindMarkers function (minpct = 0.25 and p value < 0.01). Cell identity was assigned on the basis of marker genes in each cluster (Supplementary Table 4). FindMarkers was used to calculate differentially expressed genes (DEGs) by comparing CD300e^+/+^ and CD300e^-/-^ cells within each cluster (Bonferroni adjusted p value < 0.05). Gene set enrichment analysis (GSEA) of the mouse ortholog hallmark gene sets (MH Mouse MSigDB, v2024.1. Mm) was provided with the fgsea package (Benjamini‒Hochberg adjusted p value < 0.25 v1.34.0, [47–49]) (Supplementary Table 5). The AddModuleScore from the Seurat package was used to estimate a score per cell representing the cumulative gene expression of the DEGs of the human Mono_a cluster.

All analyses were conducted in R (version 4.5.0, 2025-04-11) via RStudio.

### Statistical analyses

Statistical analyses were performed with GraphPad Prism 8.0 software via one-way or two-way analysis of variance (ANOVA) followed by Šídák’s multiple comparisons test or Tukey’s test, as appropriate. For two-group comparisons, two-tailed Student’s t tests were employed. All the data are presented as the means ± standard deviations (SDs), unless otherwise stated, and differences between groups were considered statistically significant at p<0.05. *p value≤0.05; **p value≤0.01; ***p value≤0.001; ****p value≤0.0001.

## Results

### Colorectal cancer promotes CD300e expression in dysfunctional tumor-infiltrating macrophages

To address the involvement of CD300e-expressing monocytes/macrophages in the colorectal cancer TME, we first investigated their expression in matched tumor and normal colon mucosa (NM) derived from CRC patients. Compared with matched normal tissue, tumor mucosa presented significantly increased expression of CD300e (Figure 1A), in line with previous observations linking CD300e to immunosuppressive macrophage programming and impaired antigen presentation [27]. Building on our earlier findings that TAMs in CRC downregulate MHC-II and suppress T-cell proliferation in response to tumor-derived signals [38], we assessed whether CD300e expression is associated with similar transcriptional alterations. In the same patient samples, tumor-infiltrating macrophages showed reduced MHC-II expression, accompanied by significant downregulation of CIITA and STAT1, the latter being a key transcription factor in the IFN-γ–CIITA axis (Figure 1B). These data suggest that the immunosuppressive phenotype of CD300e⁺ macrophages may result, at least in part, from impaired STAT1 signaling, leading to defective antigen presentation. Immunohistochemical analysis revealed that the increase in CD300e expression within CRC tissue reflects enhanced infiltration of CD163⁺ macrophages co-expressing CD300e. Importantly, the proportion of CD300e⁺/MHC-II^low^ macrophages was significantly elevated in tumor tissue relative to matched normal mucosa (Figure 1C), further supporting a role for CD300e in fostering a suppressive myeloid compartment within the CRC microenvironment.

**Figure 1:**
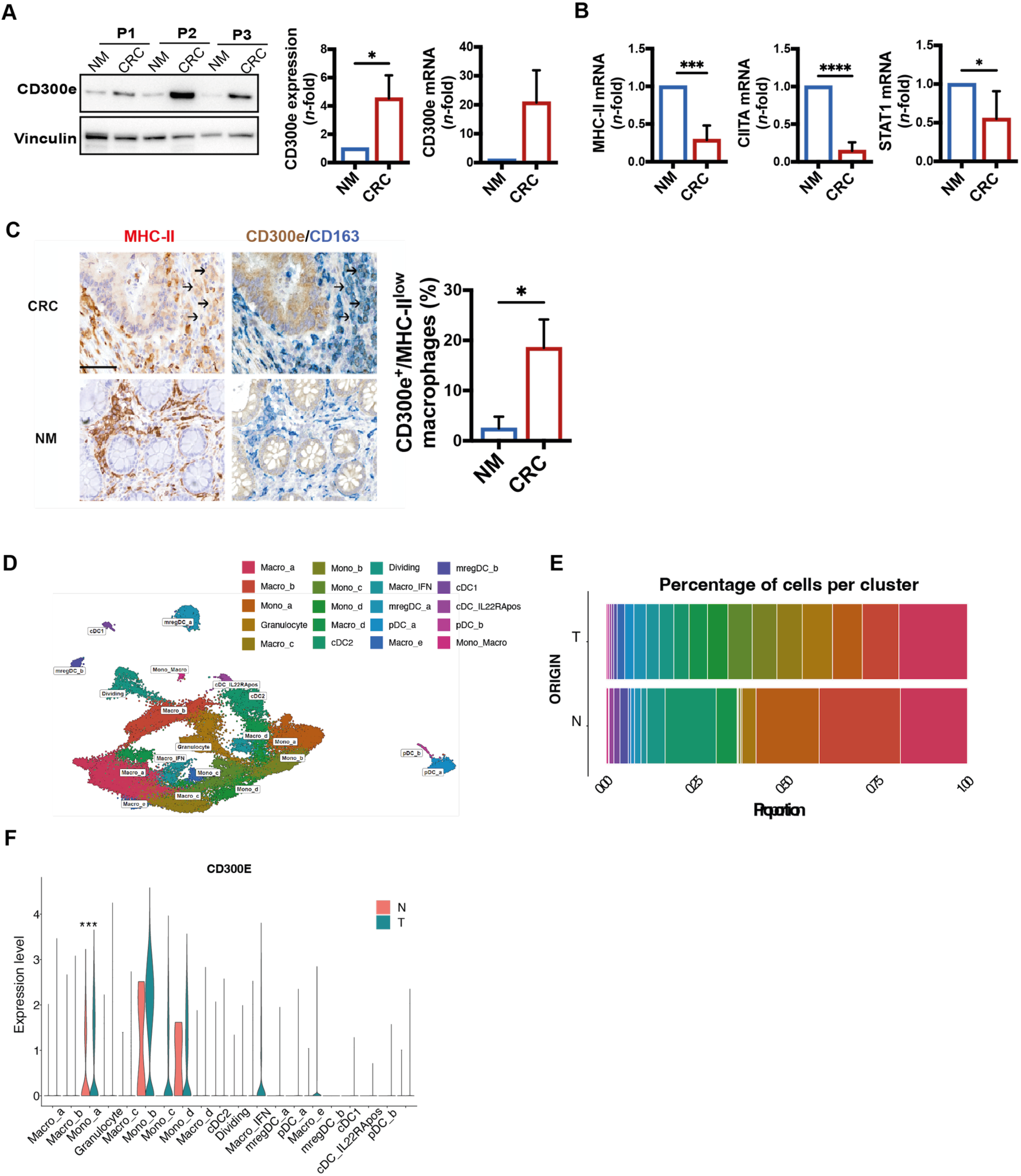
CD300e is expressed by tumor-infiltrating macrophages in colorectal cancer and is associated with the immunosuppressive profile. A) CD300e expression by western blot and qPCR in CRC tissue vs matched normal mucosa (mean ± SD of 8 CRC patients). *p value ≤0.05. B) qRT‒PCR expression of MHC-II, CIITA and STAT1 (mean ± SD of 6 CRC patients). ***p value ≤0.001; ****p value ≤0.0001. C) IHC staining of macrophages (CD163^+^ blue) expressing CD300e (brown) and MHC-II (red); arrows highlight CD300e^+^/MHC-II^low^ macrophages. The histogram shows the percentage of CD300e^+^/MHC-II^low^ macrophages expressed as the mean ± SD of 8 CRC patients. Statistical significance was determined by Student’s t test. CRC = colorectal cancer; NM = normal colonic mucosa. *p value ≤0.05. D) UMAP plot showing the transcriptional states of myeloid cells infiltrating colorectal cancer samples identified via scRNA-seq. E) Bar plot showing the contribution of each cluster identified via scRNA-seq divided by the origin of the sample. T, tumoral sample; N, tumor adjacent sample. F) Violin plot showing the normalized expression levels of CD300E across the clusters identified via scRNA-seq divided by the origin of the samples. Adjusted p value from the “Seurat” DESeq2 test. *** adjusted p value < 0.001.

To further characterize the phenotype and functional properties of CD300e^+^ myeloid cells in CRC, we leveraged a publicly available scRNA-seq dataset (GSE178341) [43] from primary tumor and adjacent normal samples from untreated CRC patients. Given that *CD300E* is predominantly expressed by myeloid cells, we focused our analysis on this compartment, and we identified 20 distinct clusters organized in granulocytes, monocytes, macrophages and dendritic cells based on their transcriptional profiles and the previous authors classification (Figure 1D and Supplementary Table 2). A comparison of cluster distributions between normal and tumor tissues revealed increased monocyte infiltration in tumor samples (Figure 1E and Supplementary Table 2). Whereas monocytes and macrophages account for approximately two-thirds of the myeloid compartment in normal tissues, their proportion is notably reduced in tumors, resulting in a more heterogeneous myeloid composition. *CD300E* expression was enriched in several monocyte clusters (Mono_a, Mono_b, Mono_C and Mono_d) and in one macrophage cluster characterized by an active interferon gene signature (Macro_IFN). Although monocytes and macrophages were the most abundant cell types in normal samples, *CD300E* expression was upregulated in tumor-infiltrating cells compared with their normal counterparts. Among these clusters, the Mono_a cluster presented significantly increased mean expression of *CD300E* in tumor tissues (Figure 1F). Given the positivity for *CD300E* of tumor derived monocyte and selected macrophage clusters, and by considering that the majority of tumor-infiltrating macrophages differentiate from recruited circulating monocytes, we performed a pseudotime trajectory analysis restricted to monocytes and macrophages, which revealed the expected progression from monocyte to macrophages (Supplementary Figure 1 A, B). Notably, as shown in Supplementary Figure 1C, the expression of *CD300E* decreased along the monocyte maturation trajectory and terminated approximately when there was a switch from Macro_IFN to Macro_c. In summary, in CRC the *CD300E* expression is largely confined to tumor-infiltrating monocytes and early monocyte-derived macrophages.

### The colorectal tumor cells drives the differentiation of immunosuppressive CD300e⁺ macrophages

Given that CD300e⁺ monocyte-derived macrophages in CRC tumors display an immunosuppressive profile, we sought to determine whether this phenotype could be directly instructed by the tumor epithelium. To this end, we established an in vitro coculture system using patient-derived tumor colon organoids (TCOs) and matched normal colon organoids (NCOs), which were incubated with peripheral blood monocytes (Figure 2A). After 5 days of coculture, monocytes exposed to both NCO and TCO acquired a macrophage-like phenotype (Figure 2B), with TCO-exposed cells showing distinct immunological features. Notably, TCO promoted strong upregulation of CD300e (Figure 2C), downregulation of MHC-II and CD86, and increased expression of the anti-inflammatory markers CD206 and CD163 relative to those in NCO-exposed cells (Figure 2D and Supplementary Figure 2). Functionally, these cells also secreted increased levels of IL-6 and IL-10, two cytokines associated with tumor-promoting inflammation (Figure 2E).

**Figure 2.**
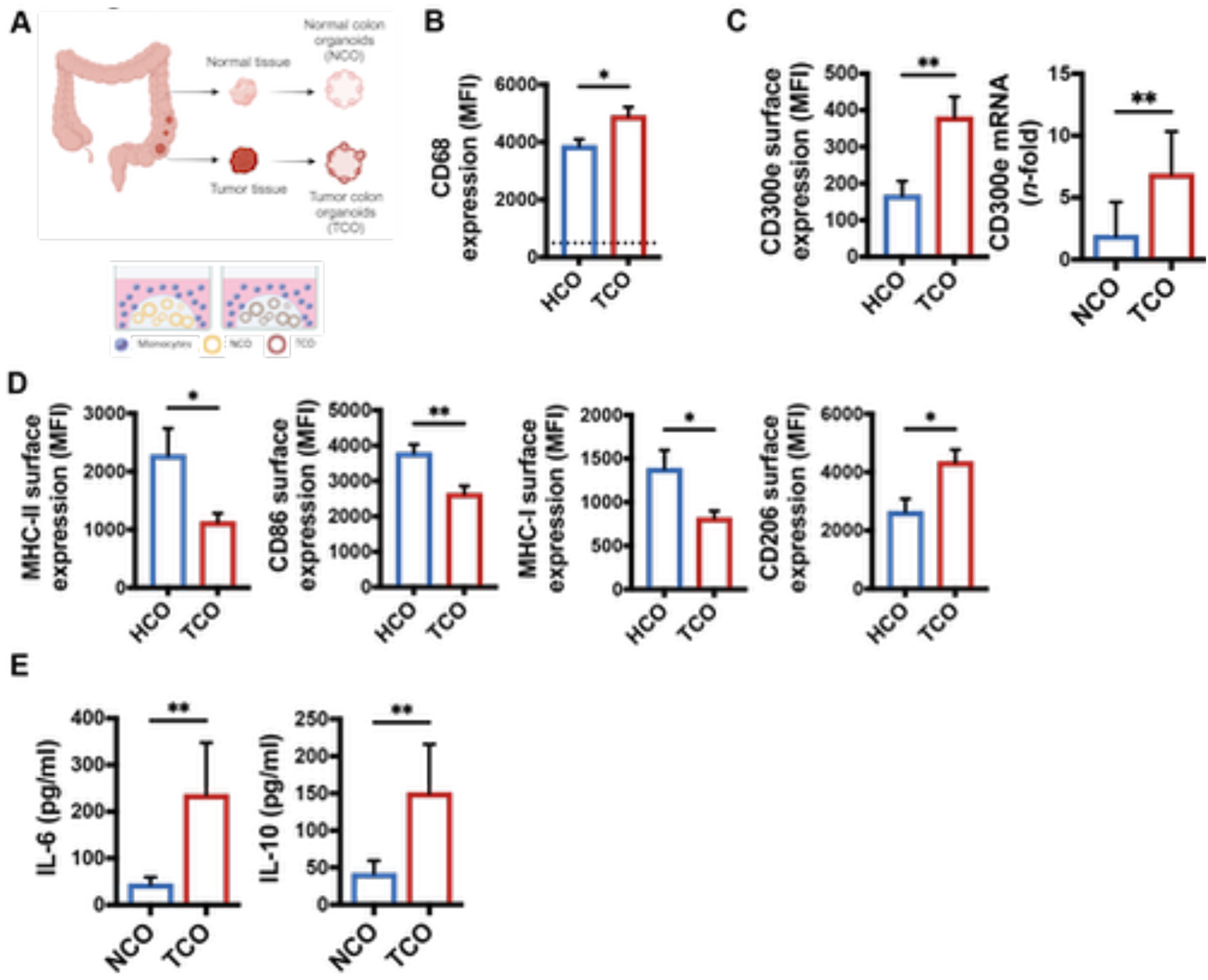
Tumor-derived colon organoids induce CD300e expression and immunosuppressive programming in human monocytes-derived macrophages. A) Scheme of the experimental set up. Image created with Biorender. NCO = normal colon organoid; TCO = tumor colon organoid. Created with Biorender. B) CD68 expression analysis by flow cytometry of monocytes at time 0 (dotted line) and monocytes-derived macrophages (MDM) cocultured for 5 d with NCO or TCO (n = 4 different monocyte donors cocultured with 4 NCO lines and 4 TCO lines). Data are expressed as Median Fluorescence Intensity (MFI). *p value ≤0.05. C) Flow cytometry and qRT-PCR of CD300e on MDMs cocultured for 5 d with NCO or TCO (n = 4 different monocyte donors cocultured with 4 NCO and 4 TCO lines). **p value ≤0.01. D) Flow cytometry analysis of MDMs (MHC-II, CD86, MHC-I, CD206) after 5 d of coculture. *p value ≤0.05; **p value ≤0.01. E) Quantification of IL-6 and IL-10 released in coculture supernatants by ELISA assay. **p value ≤0.01.

These findings indicate that the colorectal tumor epithelium actively reprograms monocytes toward CD300e^+^ macrophages with reduced antigen-presenting capacity and an immunosuppressive phenotype.

### CD300e supports colitis-associated colon cancer development

To investigate whether CD300e regulates the monocyte/macrophage phenotype, we generated a systemic *CD300E*-deficient mouse (Supplementary Figure 3A). In CD300e^+/+^ tissues, *CD300e* was expressed primarily in the blood, bone marrow (BM), and liver, which are enriched with monocytes/macrophages, and at lower levels in the heart, muscle, lymph nodes, lungs, and spleen (Supplementary Figure 3B). Efficient gene deletion was confirmed by assessing gene expression across different tissues (Supplementary Figure 3C). Notably, *CD300e* deletion did not affect leukocyte populations in the blood or BM in either sex (Supplementary Figure 3D and E), nor did it alter the phenotype of peritoneal macrophages (Supplementary Figure 3F). Histological analysis revealed no significant differences between CD300e^+/+^ and CD300e^-/-^ mice under homeostatic conditions (Supplementary Figure 3G).

To gain molecular insight into CD300e function, we performed bulk RNA-seq on BM-derived monocytes from CD300e^+/+^ and CD300e^-/-^ mice. Several genes were differentially expressed in the absence of *CD300e* (Supplementary Figure 3H). Pathway analysis revealed enrichment of genes involved in antigen presentation, immune receptor activity, cytokine signaling, and chemokine-related processes (Supplementary Figure 3I). In particular, CD300e^-/-^ monocytes presented upregulation of MHC class II genes (*H2-Ab1*, *H2-Aa*, *H2-Eb1*, and *Cd74*) and proinflammatory/immunostimulatory genes (*Tnfrsf1b*, *Il17ra*, *and Ccr7*), whereas immunosuppressive and tumor-promoting genes such as *Ccl2* [50] and *Trem2* [51] were markedly downregulated (Supplementary Figure 3J). These data suggest that CD300e shapes monocyte identity by promoting a transcriptional program associated with immunosuppression and tumor support, which is consistent with its regulatory role in human monocyte activation and function [28].

To assess whether CD300e contributes functionally to colorectal cancer development, we employed a colitis-associated colorectal cancer model using azoxymethane (AOM) and dextran sodium sulfate (DSS) [52]. Although not statistically significant, CD300e^+/+^ mice tended to have increased mortality, particularly during the first DSS cycle (Figure 3A). CD300e^-/-^ mice experienced significantly less weight loss than their CD300e^+/+^ counterparts did (Figure 3B), despite having comparable baseline weights (Supplementary Figure 4A) and similar weight gains in untreated controls (Supplementary Figure 4B). Colon shortening, a hallmark of DSS-induced injury, was significantly attenuated in CD300e^-/-^ mice (Figure 3C), accompanied by reduced intestinal permeability (Supplementary Figure 5A). The baseline colon length in the untreated mice was equivalent between the genotypes (Supplementary Figure 4C).

**Figure 3.**
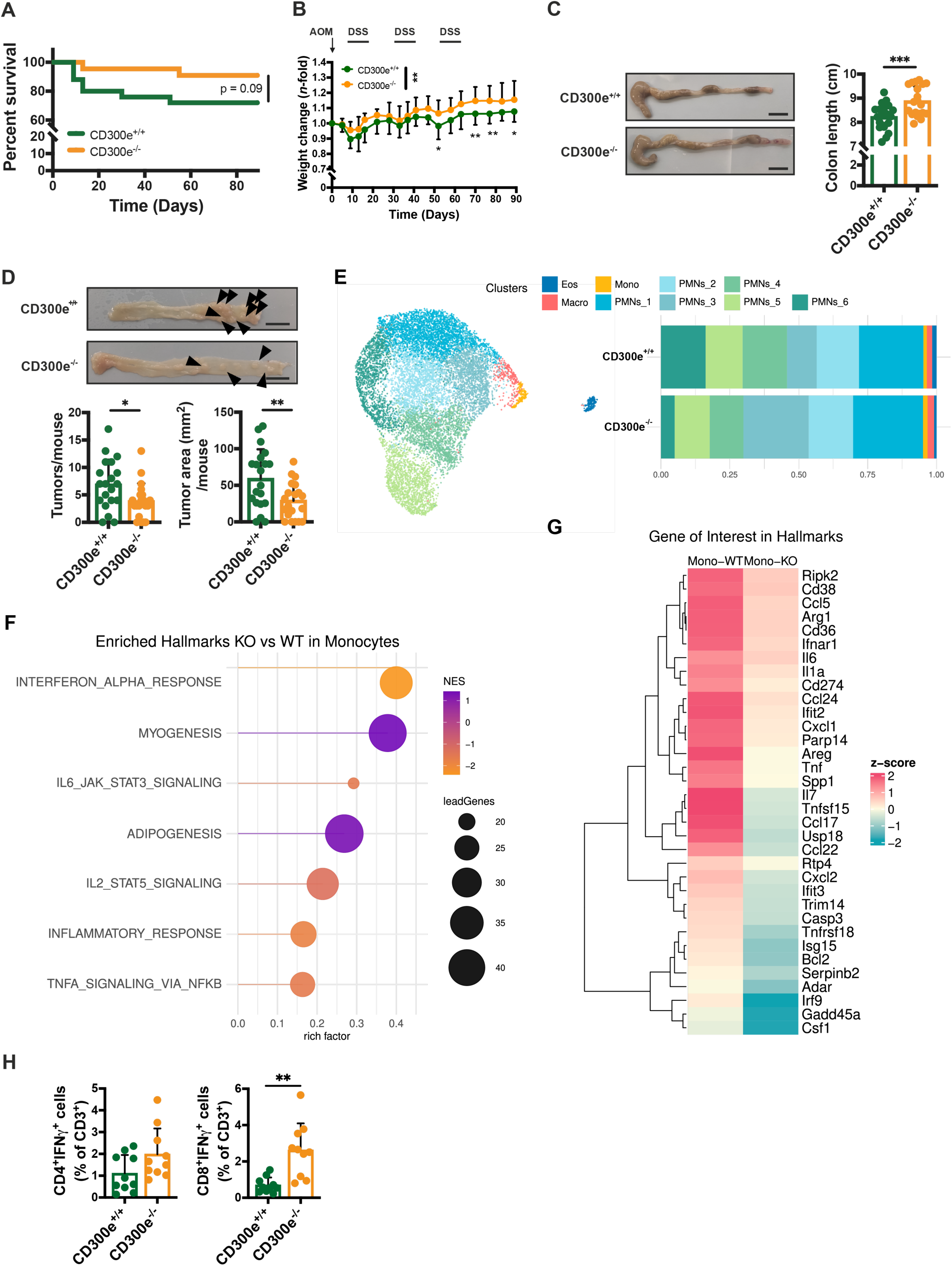
CD300e^-/-^ mice develop a lower tumor burden in the AOM/DSS CRC model. A) Kaplan‒Meier survival analysis. B) Weights of the mice throughout the AOM/DSS (n = 20 CD300e^+/+^ and 20 CD300e^-/-^ mice). Statistical significance was determined by two-way ANOVA and Šídák’s multiple comparisons test: **p value ≤0.01. C) Colon images and colon length measurements after AOM/DSS treatment (n = 20 CD300e^+/+^ and 20 CD300e^-/-^ mice; scale bar = 1 cm). ***p value ≤0.001. D) Images of longitudinally opened colons from CD300e^+/+^ and CD300e^-/-^ mice (scale bar = 1 cm). Number of tumors/mouse (left) and tumor area/mouse (right) (n = 20 CD300e^+/+^ and 20 CD300e^-/-^ mice; scale bar = 1 cm). Arrowheads indicate adenomas. *p value ≤0.05; **p value ≤0.01. E) UMAPs showing the transcriptional states of adenoma-infiltrating myeloid cells On the right, the bar plot shows the contribution of each cluster identified via scRNA-seq divided by the origin of the samples: CD300e^+/+^ and CD300e^-/-^ mice. F) Lolly plot showing the normalized enrichment scores of the GSEA of hallmarks between CD300e-/- and CD300e^+/+^. G) Heatmap of the z score of the gene of interest derived from the significant pathway from the GSEA. H) IFN-γ production by mesenteric CD4^+^ and CD8^+^ T cells. **p value ≤0.01.

Importantly, CD300e deficiency led to a significant reduction in tumor burden, as evidenced by fewer neoplastic lesions and a smaller tumor area (Figure 3D), which was confirmed histologically (Supplementary Figure 5B). Myeloid cell infiltration within adenomas was modestly reduced in CD300e^-/-^ mice, mainly due to a lower number of infiltrating neutrophils, which was consistent with the AOM/DSS model (Supplementary Figure 5C). Although monocyte/macrophage infiltration was limited overall, tumor-associated F4/80^+^ cells in CD300e^-/-^ mice presented slightly increased MHC-I and MHC-II expression (Supplementary Figure 5D). This prompted scRNA-seq analysis of adenoma- infiltrating myeloid cells to evaluate potential shifts in their transcriptional profiles.

We observed lower overall myeloid infiltration in CD300e^-/-^ mice, with neutrophils comprising the predominant population (Supplementary Table 6). Among the nine identified clusters, two represented monocytes/macrophages present at similar frequencies across genotypes, although absolute numbers were greater in CD300e^+/+^ mice (Figure 3E). A cross-species comparison revealed that murine monocytes shared a transcriptional signature with the human Mono_a cluster (Supplementary Figure 6), supporting functional conservation. Hallmark pathway analysis revealed that CD300e^-/-^ monocytes downregulated several inflammatory and tumor-associated pathways that were previously upregulated in human tumor-infiltrating Mono_a and Macro_IFN cells (Figure 3F). Overall, CD300e^+/+^ monocytes/macrophages expressed higher levels of genes associated with immunosuppression and tumor support, including *Cd36, Spp1, Il6, Cd274, Ccl5, Ccl17, Ccl22, and Ccl24* (Figure 4G and Supplementary Figure 5E) [53].

**Figure 4.**
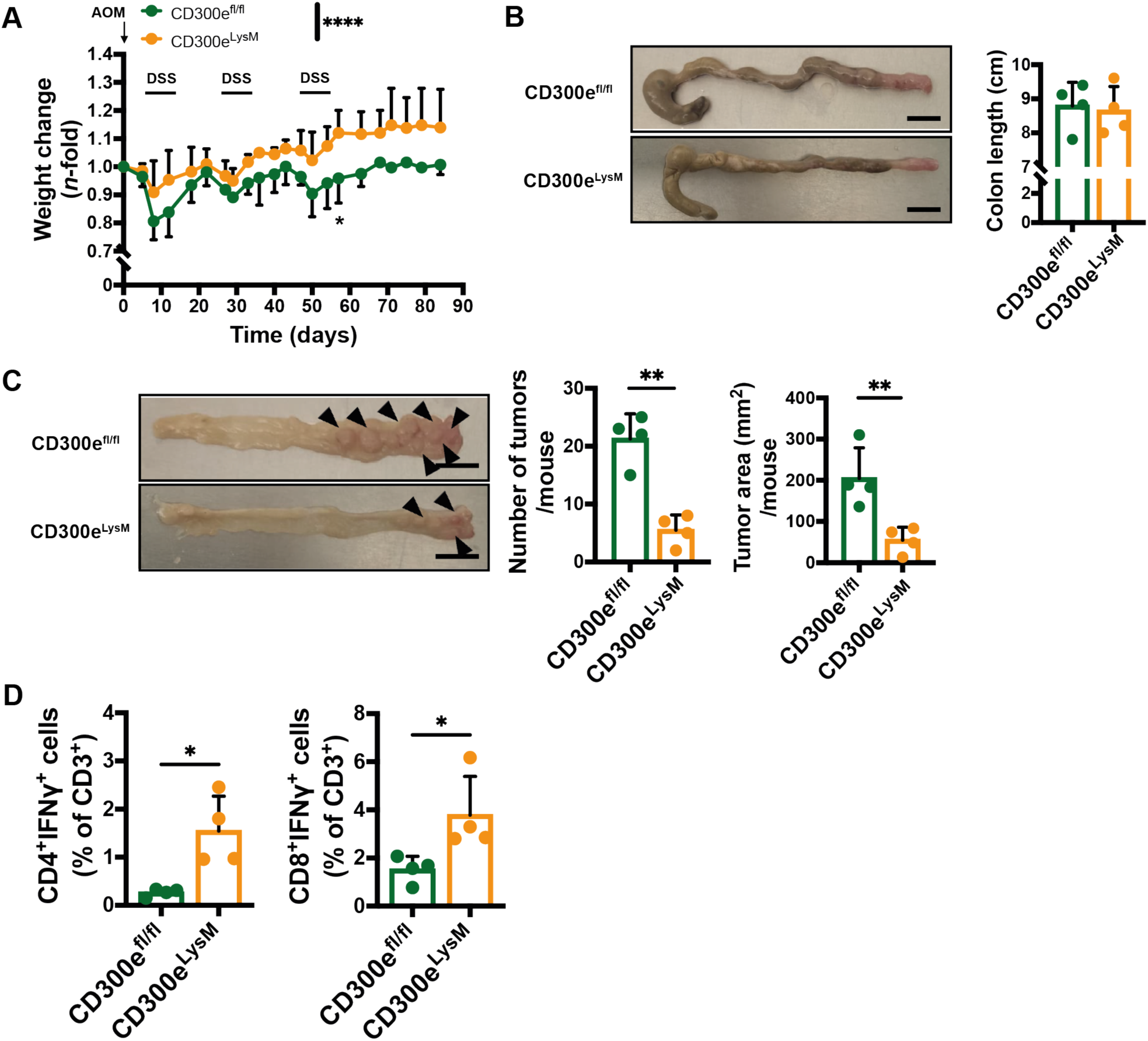
CD300e conditional knockout (CD300e^fl/fl^LysM^Cre^) mice develop a lower tumor burden. A) Weights of the mice throughout the AOM/DSS (n = 4 CD300e^fl/fl^ and 4 CD300e^LysM^ mice). Statistical significance was determined by two-way ANOVA and Šídák’s multiple comparisons test: ****p value ≤0.0001. B) Colon images and colon length measurements after AOM/DSS treatment (scale bar = 1 cm). Arrowheads indicate adenomas. C) Images of longitudinally opened colons from CD300e^+/+^ and CD300e^-/-^ mice (left; scale bar = 1 cm). Number of tumors/mouse (middle) and tumor area/mouse (right). Statistical significance was determined by Student’s t test. **p value ≤0.01. D) IFN-γ production by mesenteric CD4^+^ and CD8^+^ T cells. *p value ≤0.05.

Although tumor-infiltrating T cells were present at a low frequency, their CD4⁺ and CD8⁺ proportions were similar between genotypes (Supplementary Figure 5F). However, CD300e⁻^/^⁻ mice presented a significantly greater frequency of IFN-γ–producing CD8⁺ T cells in mesenteric lymph nodes (Figure 4H), indicating enhanced systemic antitumor immunity in the absence of CD300e. No differences in T-cell activation were detected in untreated animals (Supplementary Figure 4D–E).

To confirm that the observed effects were due to myeloid-intrinsic CD300e, we generated conditional knockout (cKO) mice in which CD300e deletion was restricted to the myeloid compartment (indicated as CD300e^LysM^) (Supplementary Figure 7A). These cKO mice recapitulated the phenotype observed in global knockouts, showing reduced weight loss, decreased tumor burden, and increased frequencies of IFN-γ–producing CD4⁺ and CD8⁺ T cells (Figure 4A–D and Supplementary Figure 7B–C).

### Lack of CD300e enhances antitumor immunity by reprogramming tumor-infiltrating myeloid cells and increasing T-cell function

To further investigate the role of CD300e in shaping the tumor immune landscape, we employed a subcutaneous CRC model by injecting the MC38 colon cancer cell line into CD300e^-/-^ and CD300e^+/+^ mice—an established model in which tumor progression is known to rely on the activity of tumor-infiltrating macrophages [54]. While MC38 tumors grew progressively in CD300e^+/+^ mice, their growth was significantly impaired in CD300e^-/-^ mice (Figure 5A, B), in line with the findings from the AOM/DSS CRC model. To characterize the immune microenvironment, we performed flow cytometry analysis of MC38 tumors 21 days postinjection (the gating strategy is shown in Supplementary Figure 8). CD11b^hi^ myeloid cells were less abundant in CD300e^-/-^ tumors because of a reduction in both neutrophils (Figure 5C) and monocytes (CD11b^+^Ly6C^+^Ly6G^-^) and monocyte-derived macrophages (CD11b^+^Ly6C^lo^F4/80^+^) (Figure 5D, F; Supplementary Figure 9). Despite their lower abundance, monocytes and macrophages infiltrating CD300e^-/-^ tumors presented significantly higher expression levels of MHC-II and MHC-I molecules (Figure 5E, G), suggesting a greater immunostimulatory profile in the absence of CD300e.

**Figure 5.**
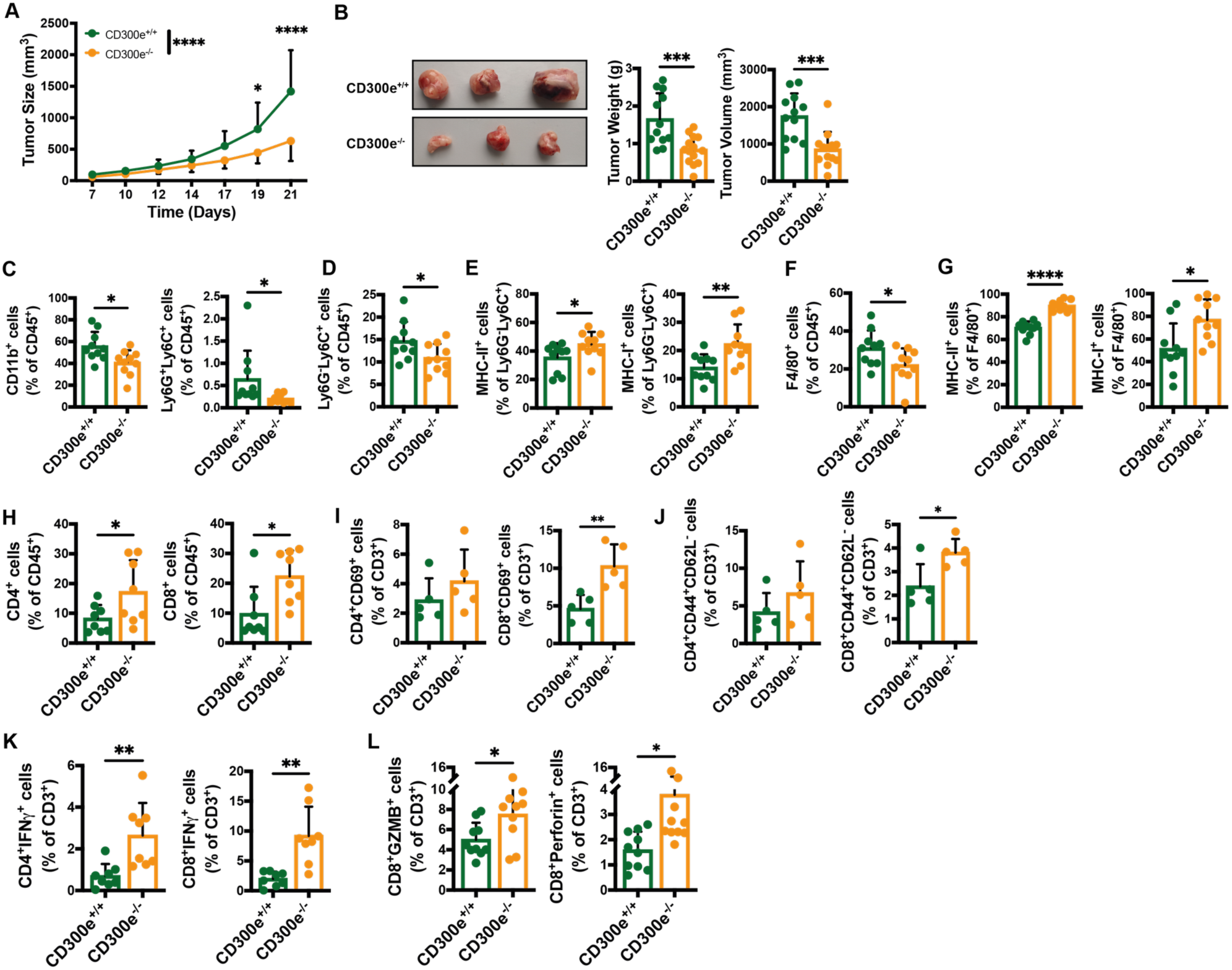
CD300e ablation delays MC38 tumor growth and modifies the tumor-immune infiltrate. A) Growth kinetics of MC38 colon adenocarcinoma in mice (mean ± SD of n = 13 CD300e^+/+^ and 13 CD300e^-/-^ mice). Statistical significance was determined by two-way ANOVA and Šídák’s multiple comparisons test: * p value ≤0.05; **** p value ≤0.0001. B) Images of tumors (left), tumor weights (middle) and final tumor volumes (right). ** p value ≤0.01. C) Myeloid cells were identified as CD45^+^CD11b^+^ and neutrophils were identified as CD45^+^CD11b^+^Ly6G^+^Ly6C^+^ and are presented as a percentage of CD45^+^ cells (n = 10 CD300e^+/+^ and 10 CD300e^-/-^ mice). * p value ≤0.05. D) Ly6G^-^Ly6C^+^ monocytes gated on CD45^+^CD11b^+^ cells are shown as a percentage of CD45^+ cells^. *p value ≤0.05. E) MHC-II- and MHC-I-expressing tumor-infiltrated monocytes gated on CD45^+^CD11b^+^Ly6G^-^Ly6C^+^ cells are shown as percentages of Ly6G^-^Ly6C^+ cells^. *p value ≤0.05; **p value ≤0.01. F) F4/80^+^ macrophages gated on CD45^+^CD11b^+^Ly6C^-^ cells are shown as a percentage of CD45^+ cells^. *p value ≤0.05. G) MHC-II- and MHC-I-expressing tumor-infiltrated macrophages gated on CD45^+^CD11b^+^Ly6C^-^ F4/80+ cells are shown as percentages of F4/80^+^ cells. *p value ≤0.05; ****p value ≤0.0001. H) CD4^+^ and CD8^+^ T lymphocyte populations in tumors gated on CD45^+^CD3^+^ cells are shown as percentages of CD45^+^ cells (means ± SDs of 8 CD300e^+/+^ and 8 CD300e^-/-^ mice). *p value ≤0.05. I) Early active CD4^+^ and CD8^+^ T cells were identified as CD69^+^ cells and are presented as the percentage of CD3^+^ cells (mean ± SD of 5 CD300e^+/+^ and 5 CD300e^-/-^ mice). **p value ≤0.01. J) Effector memory CD4^+^ and CD8^+^ T cells were identified as CD44^+^CD62L^-^ cells and are presented as percentages of CD3^+ cells^. *p value ≤0.05. K) IFN-γ production by tumor-infiltrating CD4^+^ and CD8^+^ T cells is shown as the percentage of CD3^+^ cells (mean ± SD of 8 CD300e^+/+^ and 8 CD300e^-/-^ mice). **p value ≤0.01. L) GZMB and perforin expression by tumor-infiltrating CD8^+^ T lymphocytes is shown as the percentage of CD3^+^ cells (mean ± SD of 10 CD300e^+/+^ and 10 CD300e^-/-^ mice). *p value ≤0.05.

Given that elevated expression of antigen-presenting molecules is often associated with improved T-cell activation, we next analyzed the lymphoid compartment. Compared with those from CD300e+/+ control mice, tumors from CD300e^-/-^ mice presented a greater proportion of both CD4^+^ and CD8^+^ T cells (Figure 5H; gating strategy detailed in Figure S10). In addition, CD300e^-/-^ tumors contained increased frequencies of activated and effector memory T cells within both the CD4^+^ and CD8^+^ subsets (Figure 5I, J), indicating enhanced T-cell activation. Functional profiling revealed a significant increase in the number of IFN-γ-producing CD4^+^ and CD8^+^ tumor-infiltrating T lymphocytes (TILs) in the CD300e^-/-^ mice (Figure 5K). This heightened activation was accompanied by greater cytotoxic potential, as CD8⁺ TILs from CD300e^-/-^ tumors expressed higher levels of granzyme B (GZMB) and perforin (Figure 5L).

These findings were corroborated in cKO; similar to those in full knockouts, CD300e cKO mice presented reduced tumor growth, decreased myeloid infiltration, increased MHC expression on myeloid cells, and an enhanced T-cell response characterized by elevated IFN-γ production and CD8+ cytotoxic activity (Supplementary Figure 11).

To directly compare the functional properties of macrophages from CD300e+/+ and CD300e^-/-^ mice, we performed an ex vivo phagocytosis assay on F4/80^+^ tumor-infiltrating macrophages. CD300e^-/-^ macrophages exhibited significantly greater phagocytosis capacity (Figure 6A), which is consistent with the notion that TAMs adopt a less phagocytic, immunosuppressive phenotype during tumor progression [55]. We next assessed their capacity to support T-cell activation by coculturing isolated tumor macrophages with autologous T cells labeled with CellTrace proliferation dye. Strikingly, compared with their CD300e+/+ counterparts, CD300e^-/-^ macrophages elicited significantly greater T-cell proliferation (Figure 6B).

**Figure 6:**
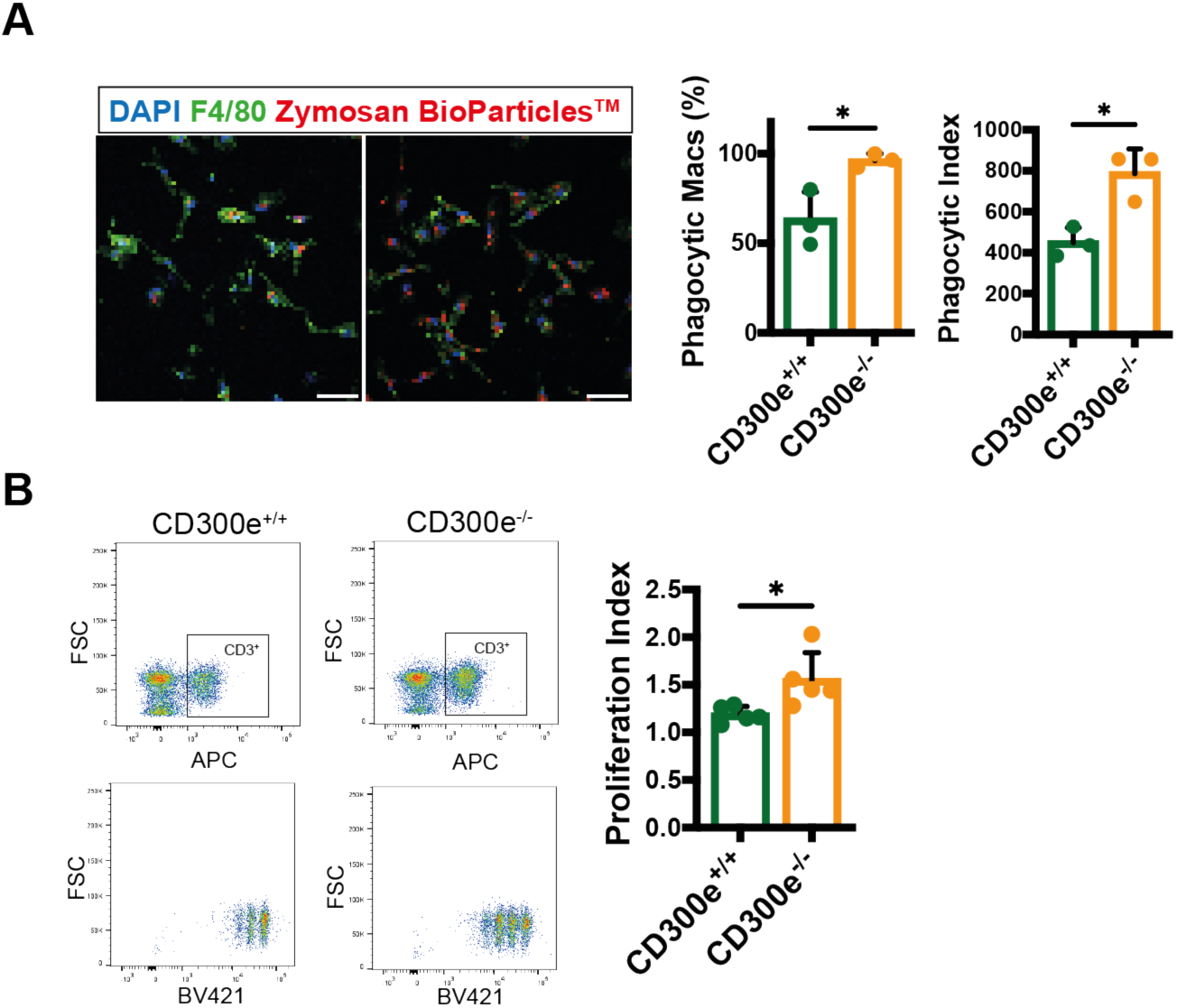
CD300e deficiency improves phagocytic capacity and T-cell proliferation. A) Representative images of the phagocytosis assay of F4/80+ macrophages isolated from CD300e^+/+^ and CD300e^-/-^ tumors incubated with Zymosan A BioParticles™. Nuclei were stained with DAPI (blue), F4/80 was used as a macrophage marker (green), and Zymosan BioParticles™ are visualized in red (left). Scale bar = 50 μm. The percentage of phagocytic macrophages (middle) was calculated as the number of phagocytic macrophages (macrophages that internalized ≥1 bioparticle) among the total number of macrophages; ten randomly acquired images per sample were analyzed (n=3 CD300e^+/+^ and 3 CD300e^-/-^ mice; right panel). The phagocytic index (right) was calculated as the percentage of phagocytic macros × the mean number of fluorescent beads per macrophage. *p value ≤0.05. B) Proliferation index of T cells as measured by CellTrace™ Violet dilution after 4 d of activation in vitro with anti-CD3 and anti-CD28 antibodies in the presence of allogenic tumor-isolated F4/80^+^ cells from CD300e^+/+^ and CD300e^-/-^ mice at a 1:3 ratio. *p value ≤0.05.

Together, these results demonstrate that CD300e deficiency promotes the acquisition of a more immunostimulatory macrophage phenotype, which in turn enhances T-cell activation, expansion, and cytotoxicity in the TME.

### CD300e expression on macrophages is sufficient to promote tumor growth and suppress the antitumor T-cell response

To determine whether the tumor-promoting effects of CD300e are directly mediated by its expression on macrophages, we performed adoptive transfer experiments using tumor-infiltrating macrophages isolated from CD300e^-/-^ and CD300e^+/+^ mice. These macrophages were co-injected at a 1:3 ratio with MC38 tumor cells into CD300e^+/+^ recipient mice (Figure 7A). Notably, tumors from mice that received CD300e^-/-^-derived macrophages exhibited significantly slower tumor growth than those from mice that received CD300e^+/+^ macrophages (Figure 7B, C). Although the overall infiltration of monocytes and macrophages within tumors was comparable between the groups, the TIL profile recapitulated the immune phenotypes observed in CD300e^-/-^ hosts. Specifically, recipient mice injected with CD300e^-/-^ macrophages presented increased frequencies of IFN-γ–producing T cells (Figure 7D) and greater cytotoxic potential among CD8⁺ lymphocytes (Figure 7E).

**Figure 7:**
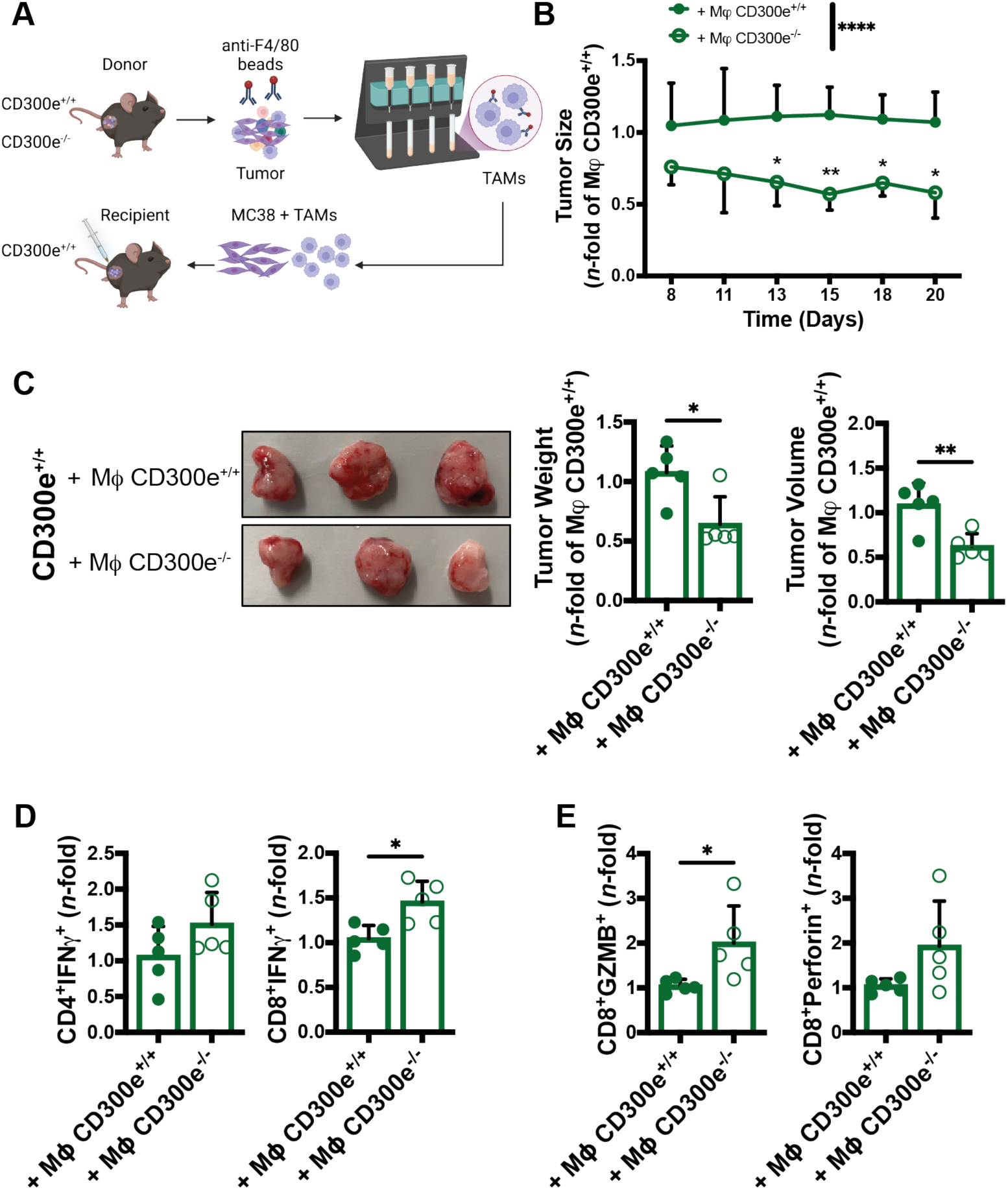
Tumor-infiltrating macrophages lacking CD300e expression are responsible for modulating the tumor microenvironment in CD300e^+/+^ mice. A) Schematic representation of the experimental workflow. Created with Biorender. B) Growth kinetics of tumors in CD300e^+/+^ mice (mean ± SD of n = 5 CD300e^+/+^ macrophages and 5 CD300e^-/-^ macrophages). Statistical significance was determined by two-way ANOVA and Šídák’s multiple comparisons test: *p value ≤0.05; **p value ≤0.01; **** p value ≤0.0001. C) Images of tumors (left), tumor weights (middle) and final tumor volumes (right). *p value ≤0.05; **p value ≤0.01. D) IFN-γ production by tumor-infiltrating CD4^+^ and CD8^+^ T cells is expressed as an n-fold difference between CD300e^+/+^ mice injected with CD300e^-/-^ macrophages and CD300e^+/+^ mice injected with CD300e^+/+^ macrophages. *p value ≤0.05. E) GZMB and perforin production by tumor-infiltrating CD8^+^ T lymphocytes expressed as *n*-fold of CD300e^+/+^ mice injected with CD300e^-/-^ macrophages vs CD300e^+/+^ mice injected with CD300e^+/+^ macrophages. *p value ≤0.05.

To confirm these findings in an independent system, we repeated the coinjection experiment using bone marrow-derived macrophages (BMDMs) polarized with IL-4 to mimic the TAM phenotype. When CD300e⁺^/^⁺ recipient mice were coinjected with MC38 cells and either CD300e⁻/⁻ or CD300e⁻^/^⁻ BMDMs at a 1:3 ratio (Supplementary Figure 12A), mice receiving CD300e⁻^/^⁻ BMDMs exhibited reduced tumor growth (Supplementary Figure 12B–C). This effect was accompanied by a greater proportion of macrophages expressing MHC-I and MHC-II (Supplementary Figure 13D) and increased infiltration of activated T cells (Supplementary Figure 12E–F).

Conversely, when tumor-isolated macrophages from CD300e^+/+^ or CD300e^-/-^ donors were injected into CD300e^-/-^ recipient mice (Supplementary Figure 13A), those receiving CD300e^+/+^ macrophages presented enhanced tumor growth (Supplementary Figure 13 B, C), coupled with reduced frequencies of IFN-γ-producing T cells and diminished CD8+ cytotoxic activity (Supplementary Figure S13 D-F).

Taken together, these data provide direct evidence that CD300e expression on macrophages is sufficient to drive tumor progression by dampening antitumor T-cell responses, highlighting its central role in orchestrating an immunosuppressive TME.

## Discussion

CD300e is a relatively understudied member of the CD300 family, originally described as an activating molecule selectively expressed by cells of the myeloid lineage [25,26]. While emerging evidence has implicated CD300e in modulating immune response during chronic inflammation and infection [27,28], its role in cancer immunity has remained unexplored.

In this study, we provide compelling evidence that CD300e plays an active and previously unrecognized role in CRC progression by enforcing an immunosuppressive program in TAMs and limiting antitumor T-cell immunity. Specifically, we show that CD300e is selectively expressed by tumor-infiltrating monocytes and monocyte-derived macrophages in human CRC, especially within a subset exhibiting IFN-driven transcriptional features.

Interestingly, despite the enrichment of IFN-related gene signature, these CD300e^+^ macrophages exhibit marked downregulation of key IFN effectors, including STAT1 and CIITA, the latter being the master regulator of MHC-II transcription. This paradoxical finding is consistent with our previous observations [28], and suggests dysregulated or suppressed IFNγ response within the tumor microenvironment, leading to impaired antigen-presentation and contributing to local immune suppression.

Importantly, our data position CD300e not merely as a marker of immunosuppressive TAMs, but as a functional regulator that acts upstream of the transcriptional downregulation of the STAT1–CIITA axis. This mechanistic link provides a functional explanation for earlier observations in CRC [38], where MHC-II^low^/CD206^high^ TAMs failed to support effective T-cell activation.

Building on prior findings that CD300e engagement can suppress MHC-II expression and macrophage-mediated T-cell proliferation [27], our study now offers in vivo validation of this immunoregulatory pathway in the context of cancer. By identifying CD300e as a central modulator of TAM dysfunction, we add a new layer of insight into the molecular mechanisms by which the tumor microenvironment subverts innate immune cells to escape immune surveillance.

Using a coculture system of patient-derived tumor colon organoids and primary human monocytes, we further demonstrated that the tumor epithelium can directly induce CD300e expression and reprogram macrophages toward an immunosuppressive phenotype. These tumor-educated macrophages exhibited hallmark features of immune regulation, including downregulation of MHC-II and CD86, upregulation of CD206 and CD163, and increased secretion of IL-6 and IL-10. These data suggest that tumor-derived cues are sufficient to drive CD300e expression and promote the emergence of an immunoregulatory TAM phenotype, capable of dampening antitumor immunity. In vivo, the functional relevance of CD300e was confirmed in two complementary models of CRC: the AOM/DSS-induced colitis-associated cancer model and the MC38 subcutaneous tumor model. Genetic ablation of CD300e in these settings resulted in a marked reduction of tumor burden, preserved intestinal barrier function, and enhanced antitumor immunity both systemically and within the tumor microenvironment. CD300e-deficient mice showed decreased infiltration of protumor myeloid cells, increased expression of MHC-I and MHC-II molecules both on monocytes and monocyte-derived macrophages, and robust activation of CD4⁺ and CD8⁺ T cells. T cells included an increased frequency of IFN-γ-producing and cytotoxic T lymphocytes.

Functional analyses further revealed that CD300e-deficient tumor-infiltrating macrophages had increased phagocytic capacity and better promoted T-cell proliferation ex vivo, indicating enhanced macrophage effector functions and antigen-presenting capability. These features are hallmarks of macrophages that support productive antitumor immunity [56]. Together, these data provide strong evidence that CD300e suppresses macrophage effector functions and supports the establishment of a tolerogenic, immune-excluded tumor microenvironment. Our findings not only reinforce the immunosuppressive nature of CD300e-expressing TAMs but also highlight CD300e as a central node through which tumor-derived signals impair macrophage-mediated immune surveillance. By attenuating antigen presentation even in the context of an active IFN response, CD300e may serve as a critical checkpoint limiting the full activation of antitumor immunity in CRC. Conditional deletion of CD300e in myeloid cells recapitulated the antitumor effects observed in full knockout mice, and adoptive transfer experiments confirmed that CD300e expression on macrophages alone is sufficient to drive tumor growth and suppress T-cell activation. These findings firmly establish CD300e as a central regulator of macrophage-mediated immune evasion in CRC. Our data position CD300e has a central modulator of monocyte-derived macrophage function in the tumor microenvironment. While high macrophage infiltration has traditionally been associated with favorable outcomes in CRC [19], but a growing body of evidence underscore the heterogeneity and plasticity of TAMs. Notably, in CRC, high density of TAMs are now correlated with a poor prognosis [23]. Our findings add functional depth to this perspective by showing that is not the number, but rather the phenotype of TAMs that dictate their impact on tumor progression. In CD300e-deficient tumors, despite a marked reduction in macrophage infiltration, the remaining cells were functionally reprogrammed to support robust antitumor immunity. This highlights CD300e not only as a marker of suppressive TAMs but also as a driver of their tumor-promoting behavior.

In conclusion, our study identified CD300e as a previously unrecognized orchestrator of immune suppression in CRC, acting through reprogramming of TAMs. Targeting CD300e offers a promising strategy to convert these cells into active effector cell in antitumor immunity. By restoring their responsiveness to IFNγ and enhancing antigen presentation, CD300e blockade could enhance the efficacy of immunotherapy. Future studies should focus on the development of selective CD300e inhibitors and their integration into existing immunotherapy regimens, particularly in combination with immune checkpoint blockade, to overcome resistance and improve outcomes for patients with CRC.

## Availability of Data and Material

The dataset(s) supporting the conclusions of this article is (are) available in the [repository name] repository, [unique persistent identifier and hyperlink to dataset(s) in http://format].

## Author contributions

G.C. conceptually planned and supervised the study. G.C., A.B. and S.V. designed the experiments. A.B. and S.V. performed most of the experiments unless otherwise specified. G.C., A.B., S.V., L.M. S.L., M.B., S.S., M.L. and E.C. analyzed the data. S.C., S.G. and S.P. participated in the animal studies. A.B., S.V., S.G., and S.C. participated in the molecular and cellular biological experiments. S.L., M.B., and W.V. helped with the pathology-related studies. G.C., A.B., M.L., E.C. and S.S. wrote the manuscript. G.S., M.F. and W.V. participated in the clinical sample and information collection. G.S. conceptualized the clinical aspects and contributed to the clinical sample collection and data analysis. All the authors have read and approved the manuscript.

## Supporting information

Supplementary figures are included in the present version of the manuscript.

## Acknowledgment

This work was supported by AIRC MFAG 23192 (to G.C.) and by AIRC IG 29244 (to S.S.) We thank Prof. Kuo from Stanford Junior University, Redwood City, CA, USA, for kindly providing Rspo-1 cells and Prof. den Hertog from Hubrecht Institute, Utrecht, The Netherlands, for kindly providing Noggin- and WNT-producing cells. We thank Prof. Bronte from the Veneto Institute of Oncology, Padua, Italy, and Prof. Marigo from the Department of Surgical, Oncological and Gastroenterological Sciences (DiSCOG), University of Padua, Padua, Italy, for kindly providing the MC38 cell line.

We thank Javier Martin Gonzalez and Rafael Oliveira Brandão from the Transgenic Core Facility of the University of Copenhagen for the generation of the CD300e KO mouse.

