## Supplemental Figures for "CD300e is a driver of the immunosuppressive tumor microenvironment and colorectal cancer progression via macrophage reprogramming"

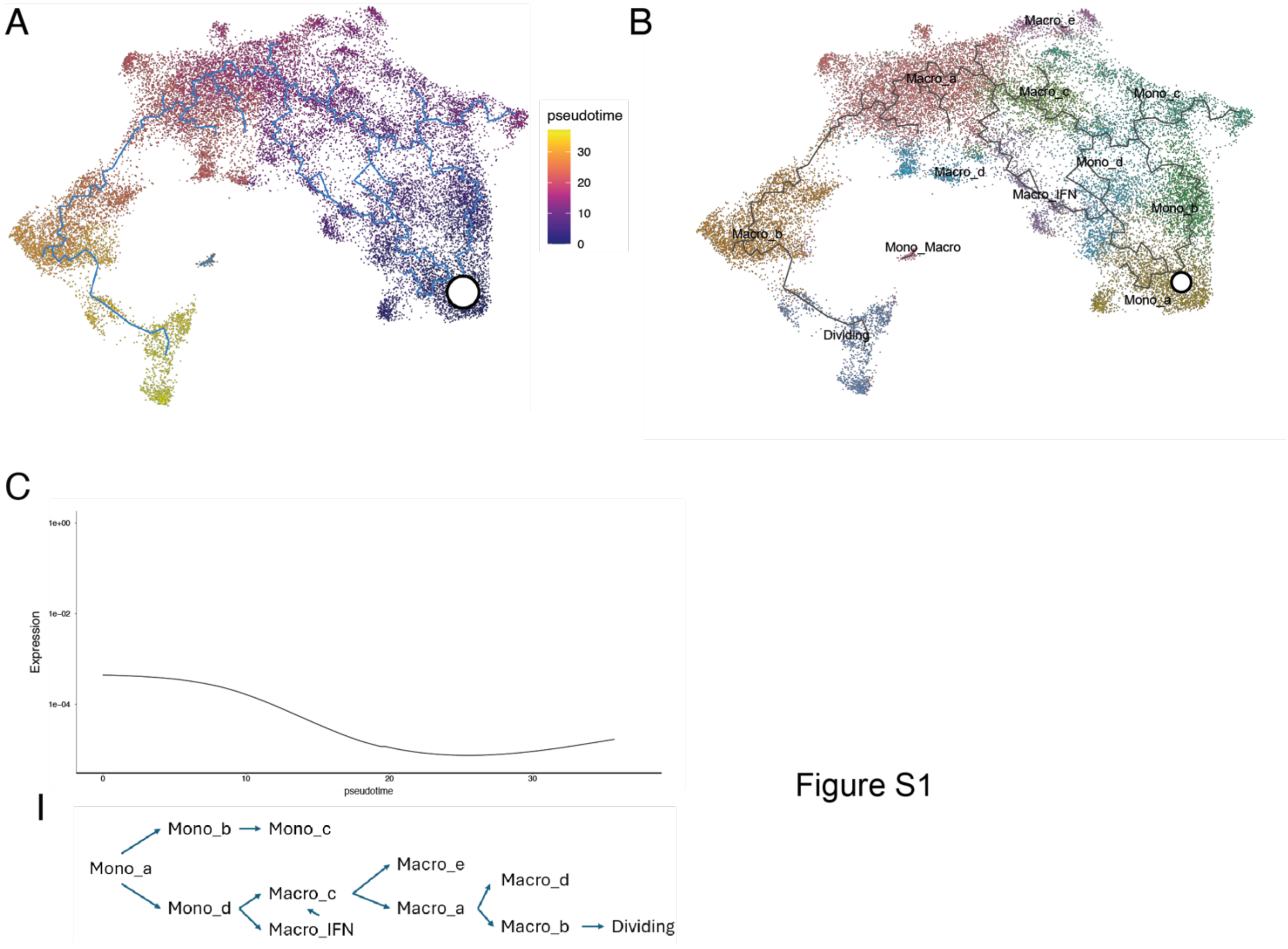


Figure S1

**Supplementary Figure 1: CD300E is a marker for tumor infiltrating monocytes and monocyte-derived macrophages.**

The trajectory inferred by the pseudo-time analysis is plotted into the UMAP devoid of dendritic cells and PMNs. The trajectory begins in the white dot, identifiable as the Mono_a cluster.

A) CD300E expression levels across the trajectory.
B) The color code shows the monocyte and macrophages cluster distribution across the UMAP and trajectory.
C) Schematic of the trajectory inferred by the pseudotime analysis.

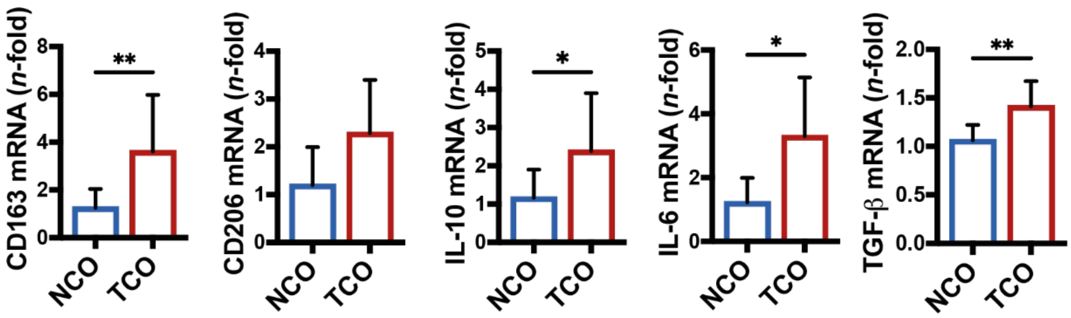


Figure S2

**Supplementary Figure 2: CD300e associates with macrophages immunosuppressive profile.**
qRT-PCR analysis of CD300e on MDM co-cultured for 5 d with NCO or TCO (n = 4 different monocyte donors co-cultured with 4 NCO and 4 TCO lines). *p-val≤0.05; **p-val≤0.01.

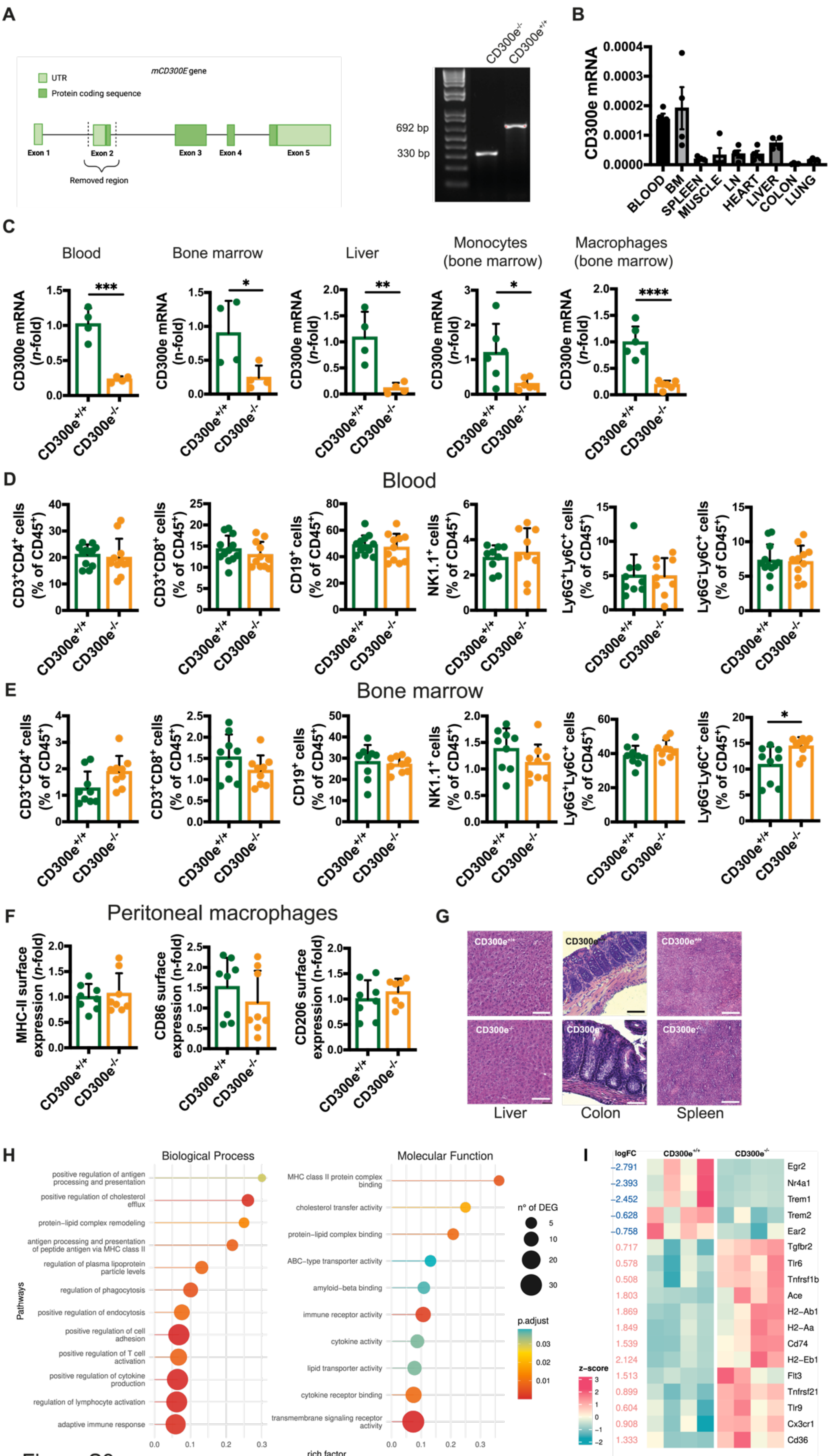
A
mCD300E gene
UTR
Protein coding sequence
Exon 1
Exon 2
Exon 3
Exon 4
Exon 5
Removed region
692 bp
330 bp
B
CD300e mRNA
BLOOD
BM
SPLEEN
MUSCLE
LN
HEART
LIVER
COLON
LUNG
C
Blood
Bone marrow
Liver
Monocytes (bone marrow)
Macrophages (bone marrow)
CD300e mRNA (n-fold)
D
Blood
CD3+CD4+ cells (% of CD45+)
CD3+CD8+ cells (% of CD45+)
CD19+ cells (% of CD45+)
NK1.1+ cells (% of CD45+)
Ly6G+Ly6C+ cells (% of CD45+)
Ly6G-Ly6C+ cells (% of CD45+)
E
Bone marrow
F
Peritoneal macrophages
MHC-II surface expression (n-fold)
CD86 surface expression (n-fold)
CD206 surface expression (n-fold)
G
Liver
Colon
Spleen
H
Biological Process
Molecular Function
Pathways
rich factor
n° of DEG
p.adjust
positive regulation of antigen processing and presentation
positive regulation of cholesterol efflux
protein-lipid complex remodeling
antigen processing and presentation of peptide antigen via MHC class II
regulation of plasma lipoprotein particle levels
regulation of phagocytosis
positive regulation of endocytosis
positive regulation of cell adhesion
positive regulation of T cell activation
positive regulation of cytokine production
regulation of lymphocyte activation
adaptive immune response
MHC class II protein complex binding
cholesterol transfer activity
protein-lipid complex binding
ABC-type transporter activity
amyloid-beta binding
immune receptor activity
cytokine activity
lipid transporter activity
cytokine receptor binding
transmembrane signaling receptor activity
I
logFC
z-score
Egr2
Nr4a1
Trem1
Trem2
Ear2
Tgfbr2
Tlr6
Tnfrsf1b
Ace
H2-Ab1
H2-Aa
Cd74
H2-Eb1
Flt3
Tnfrsf21
Tlr9
Cx3cr1
Cd36

Figure S3

**Supplementary Figure 3: CD300e$^{-/-}$ mouse characterization.**

A) Left: scheme of mouse CD300e gene with intronic (horizontal black line) and exonic (green boxes, exons 1 to 5) regions. Light green: untranslated (UTR) regions; dark green: protein-coding regions. CD300e$^{-/-}$ mice carry a deletion of 362 bp encompassing exon 2, together with upstream and downstream intronic regions. Right: representative electrophoresis gel of the genotyping of a CD300e$^{+/+}$ and a CD300e$^{-/-}$ mouse. The band in the first lane represents the deleted allele (330 bp), whereas the band in the second lane represents wild type allele (692 bp).

B) mRNA expression of CD300e was evaluated by qRT-PCR in the blood, bone marrow (BM), liver, heart, muscle, lymph nodes, lungs, spleen, and colon of CD300e$^{+/+}$ mice. Data were normalized on β-actin and 18S and as endogenous reference genes and are expressed as 2-ΔCt (mean ± SD of 4 mice).

C) mRNA expression of CD300e was evaluated by qRT-PCR in blood, BM, liver, monocytes purified from BM and BM-derived macrophages of CD300e$^{+/+}$ and CD300e$^{-/-}$ mice. Data were normalized on β-actin and 18S as endogenous reference genes and are expressed as $2^{-\Delta\Delta Ct}$ relative to CD300e$^{+/+}$ mice (mean ± SD, n = 4 CD300e$^{+/+}$ and 4 CD300e$^{-/-}$ mice for blood, BM and liver, n = 6 CD300e$^{+/+}$ and 6 CD300e$^{-/-}$ mice for monocytes and macrophages.

D, E) The percentage of CD3$^{+}$CD8$^{+}$ (T cytotoxic cells), CD3$^{+}$CD4$^{+}$ (T helper cells), CD19$^{+}$ (B cells), NK1.1$^{+}$ (natural killer cells), Ly6G$^{+}$Ly6C$^{+}$ (neutrophils) and Ly6G$^{-}$Ly6C$^{+}$ (monocytes) cells was evaluated by flow cytometry on CD45$^{+}$ blood leukocytes (D) and bone marrow leukocytes

E) in CD300e$^{+/+}$ and CD300e$^{-/-}$ mice of 8-10 weeks of age. Data are expressed as cell percentage (%) on CD45$^{+}$ leukocytes (mean ± SD of 9 CD300e$^{+/+}$ and 9 CD300e$^{-/-}$ mice).

F) The surface expression of MHC-II, CD86 and CD206 was evaluated by flow cytometry on macrophages isolated from the peritoneum. Data are expressed as *n*-fold vs macrophages from CD300e$^{+/+}$ mice ± SD of 8 CD300e$^{+/+}$ and 8 CD300e$^{-/-}$ mice.

G) Representative images of hematoxilin and eosin staining of the liver, colon and spleen of CD300e$^{+/+}$ and CD300e$^{-/-}$ mice. Scale bar = 100 μm.

H) Lolliplot graphs representing the pathways that were found most significantly deregulated in CD300e$^{-/-}$ monocytes compared to CD300e$^{+/+}$ according to the Gene Ontology (class Molecular function and Biological processes) of differentially expressed genes (DEG).

I) Heatmap showing the most differentially expressed genes between CD300e$^{+/+}$ and CD300e$^{-/-}$ monocytes, with log-fold-change for each gene shown on the left side of the plot. Statistical significance was determined by Student's t test. BM = bone marrow; LN = lymph nodes; UTR = untranslated regions.

Statistical significance was determined by Student's t test. *p-val≤0.05; **p-val≤0.01; ***p-val≤0.001; ****p-val≤0.0001.

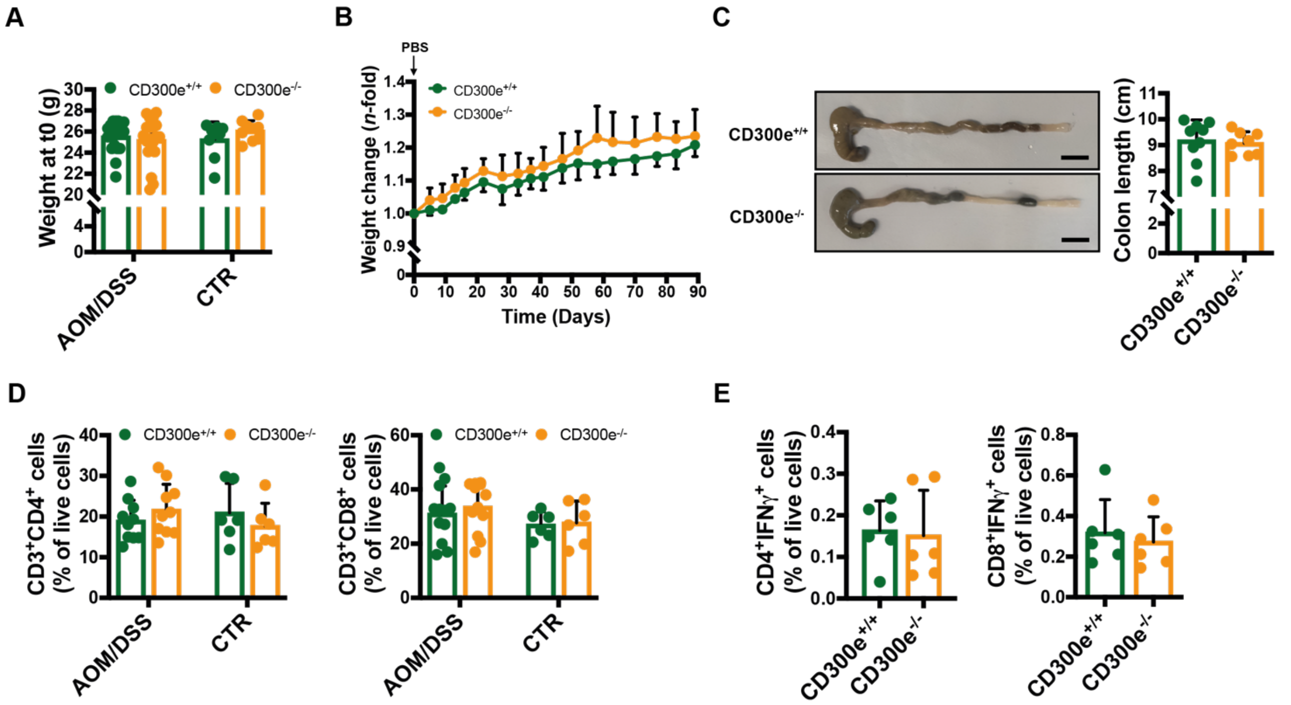


Figure S4

**Supplementary Figure 4: CD300e$^{+/+}$ and CD300e$^{-/-}$ controls to AOM/DSS treatment.**

A) Mouse weight was measured at t0, both in AOM/DSS-treated (AOM/DSS) and control (CTR) mice. Histograms represent mouse weights expressed as mean ± SD of 20 CD300e$^{+/+}$ and 20 CD300e$^{-/-}$ AOM/DSS treated mice, and 9 CD300e$^{+/+}$ and 8 CD300e$^{-/-}$ control mice.

B) The weight of control mice was recorded throughout the experimental period. Mice weights at all time points are expressed as n-fold of the weight at the beginning of the experiment (day 0, before PBS injection), ± SD (n = 9 CD300e$^{+/+}$ and 8 CD300e$^{-/-}$ mice).

C) Colon length was measured after sacrifice in control mice. Representative pictures of colons of CD300e$^{+/+}$ and CD300e$^{-/-}$ mice are shown (scale bar = 1 cm). Histogram represents colon lengths expressed as mean ± SD of 9 CD300e$^{+/+}$ and 8 CD300e$^{-/-}$ mice.

D) The percentage of CD3$^+$CD8$^+$ and CD3$^+$CD4$^+$ lymphocytes was evaluated in mesenteric lymph nodes collected from AOM/DSS-treated and control mice. Data are expressed as cell percentage (%) on live cells (mean ± SD of 10 CD300e$^{+/+}$ and 10 CD300e$^{-/-}$ AOM/DSS treated mice, 6 CD300e$^{+/+}$ and 6 CD300e$^{-/-}$ control mice).

E) The production of IFN-γ was evaluated in mesenteric lymph nodes of control mice. After overnight activation of mesenteric lymphocytes, IFN-γ levels were measured by flow cytometry. Data are expressed as percentage of IFN-γ-positive CD4$^+$ (left) and CD8$^+$ (right) T lymphocytes (mean ± SD of 6 CD300e$^{+/+}$ and 6 CD300e$^{-/-}$ mice). AOM = axozymethane; CTR = control mice; DSS = dextran sodium sulfate.

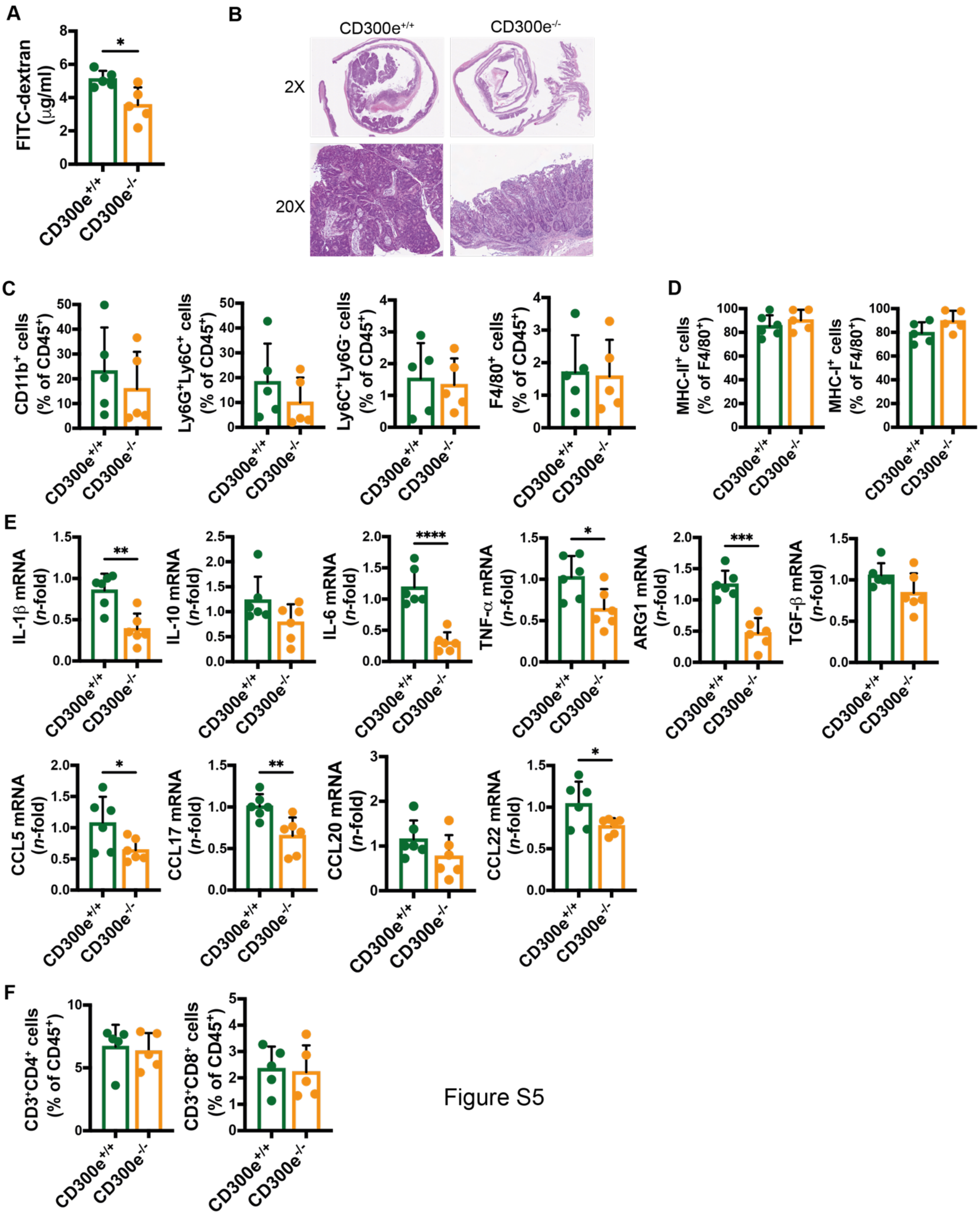
A
FITC-dextran (μg/ml)
*
CD300e+/+
CD300e-/-
B
CD300e+/+
CD300e-/-
2X
20X
C
CD11b+ cells (% of CD45+)
Ly6G+Ly6C+ cells (% of CD45+)
Ly6C+Ly6G- cells (% of CD45+)
F4/80+ cells (% of CD45+)
D
MHC-II+ cells (% of F4/80+)
MHC-I+ cells (% of F4/80+)
E
IL-1β mRNA (n-fold)
**
IL-10 mRNA (n-fold)
IL-6 mRNA (n-fold)
****
TNF-α mRNA (n-fold)
*
ARG1 mRNA (n-fold)
***
TGF-β mRNA (n-fold)
CCL5 mRNA (n-fold)
*
CCL17 mRNA (n-fold)
**
CCL20 mRNA (n-fold)
CCL22 mRNA (n-fold)
*
F
CD3+CD4+ cells (% of CD45+)
CD3+CD8+ cells (% of CD45+)

Figure S5

**Supplementary Figure 5: CD300e$^{-/-}$ mice display an improved anti-tumor immune response upon AOM/DSS treatment.**

A) After o.n. fasting, mice were gavaged with 600 mg/kg FITC-Dextran. After 3 hours, blood was collected and centrifuged to obtain plasma, and FITC fluorescence was measured. Data are expressed as mean ± SD. n=5 CD300$^{+/+}$ and 5 CD300e$^{-/-}$ mice.

B) Paraffin-embedded sections of the colon using Swiss roll technique were H&E stained. Images were obtained with ×2 (left) and ×20 (right) magnification respectively.

C) CD11b$^{+}$ myeloid cells (gated on CD45$^{+}$CD3$^{-}$CD19-NK1.1$^{-}$ cells) were identified as Ly6G$^{+}$Ly6C$^{+}$ neutrophils, Ly6G$^{-}$Ly6C$^{+}$ monocytes and F4/80$^{+}$ macrophages. Data are expressed as percentage of CD45$^{+}$ cells (mean ± SD of 5 CD300e$^{+/+}$ and 5 CD300e$^{-/-}$ mice).

D) MHC-II and MHC-I-expressing tumor-infiltrated macrophages. Data are expressed as mean percentage of F4/80+ cells ± SD.

E) qRT-PCR analysis of pro- and anti-inflammatory genes.

F) CD3$^{+}$CD4$^{+}$ and CD3$^{+}$CD8$^{+}$ T cells. Data are expressed as mean percentage of CD45$^{+}$ cells ± SD.

Statistical significance was determined by Student's t test. *p-val≤0.05; **p-val≤0.01; ***p-val≤0.001; ****p-val≤0.0001.

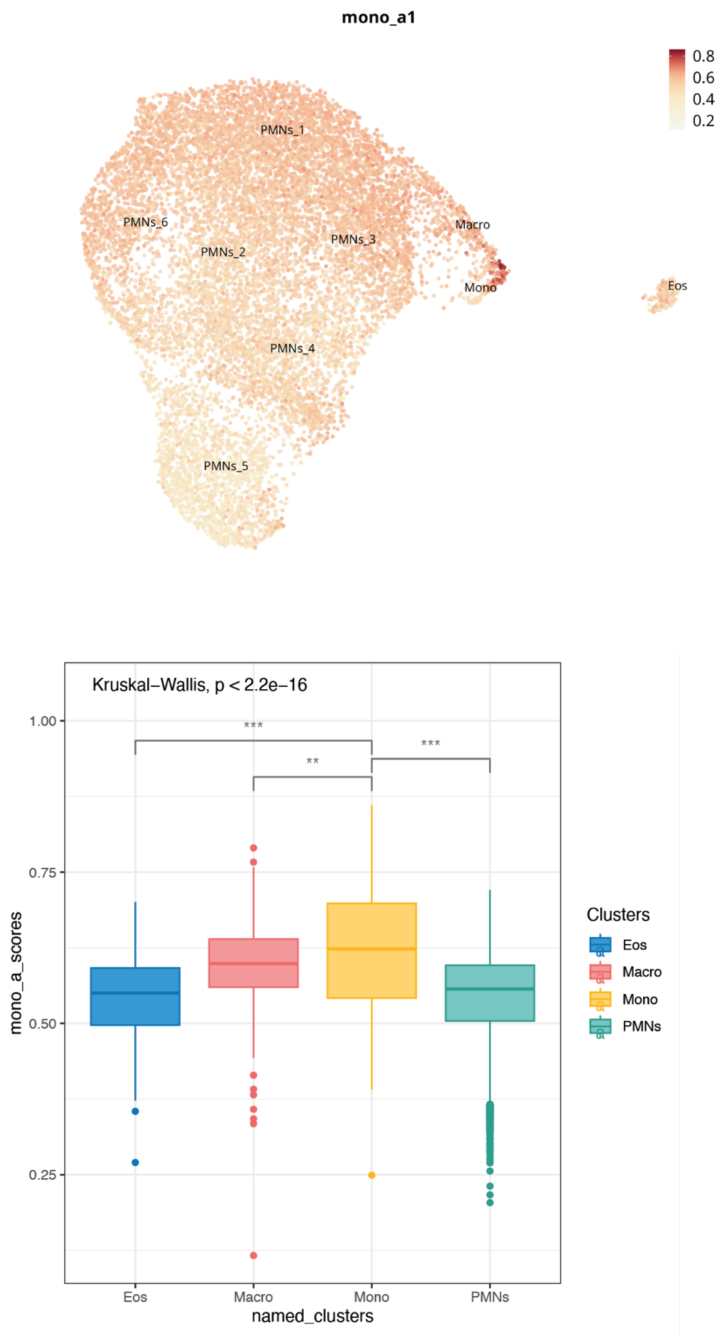


Figure S6

**Supplementary Figure S6.**

UMAP on single-cell mouse data showing the score per cell of the cumulative gene expression of the DEGs of the human Mono_a cluster (above). Box plot of the average expression levels of mono_a scores on cells in the different clusters. The Kruskal-Wallis non-parametric test was applied to compare the cluster groups.

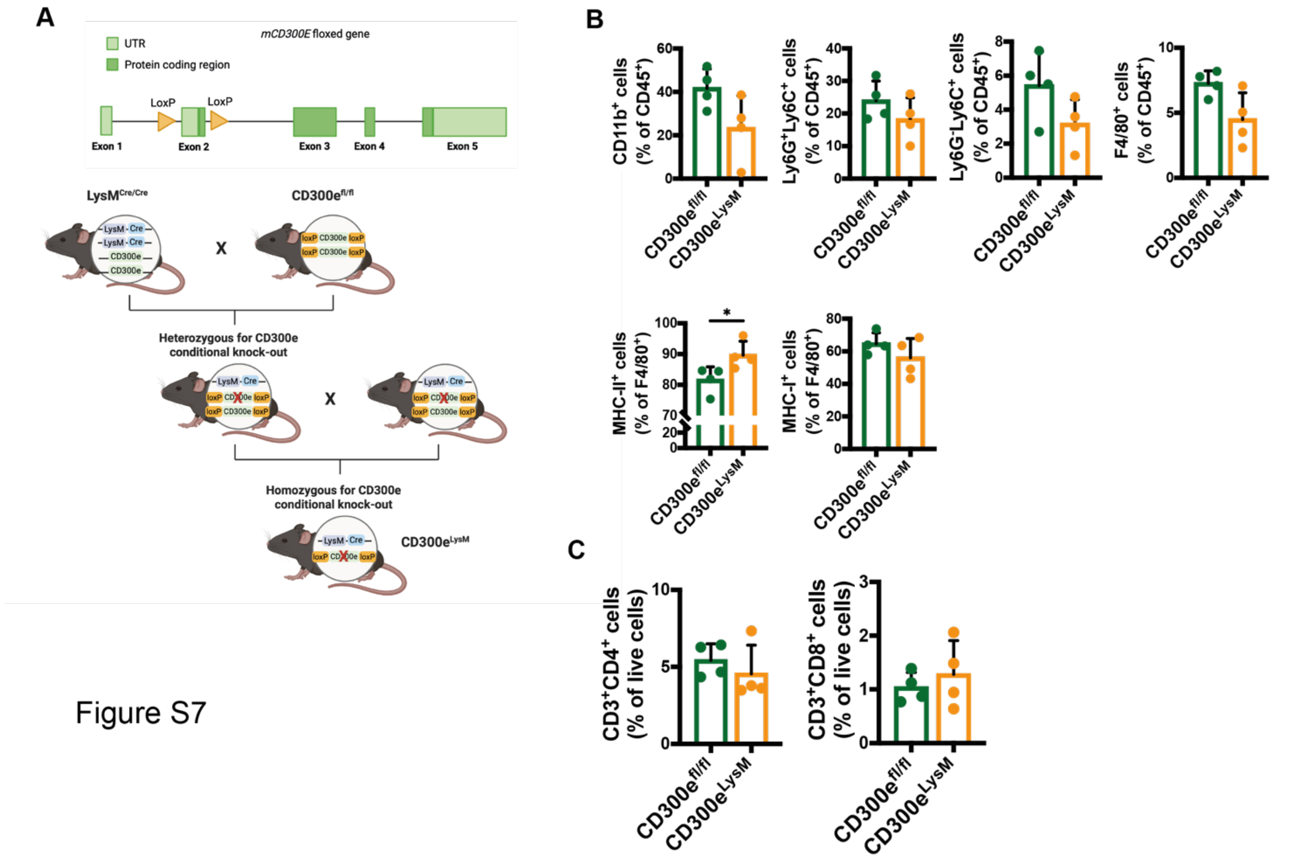


Figure S7

**Supplementary Figure 7: CD300e deletion on myeloid cells reduces tumor burden.**

A) Scheme of CD300e$^{LysM}$ mouse generation, showing loxP sites insertion flanking exon 2 and crossing with LysM Cre mouse line.

B) CD11b$^{+}$ myeloid cells (gated on CD45$^{+}$CD3$^{-}$CD19-NK1.1$^{-}$ cells) were identified as Ly6G$^{+}$Ly6C$^{+}$ neutrophils, Ly6G$^{-}$Ly6C$^{+}$ monocytes and Ly6C$^{lo}$F4/80$^{+}$ macrophages. Data are expressed as percentage of CD45$^{+}$ cells (mean ± SD of 5 CD300e$^{+/+}$ and 5 CD300e$^{-/-}$ mice). MHC-II and MHC-I-expressing tumor-infiltrated macrophages. Data are expressed as mean percentage of F4/80+ cells ± SD.

C) CD3$^{+}$CD4$^{+}$ and CD3$^{+}$CD8$^{+}$ T cells. Data are expressed as mean percentage of CD45$^{+}$ cells ± SD.

Statistical significance was determined by Student's t test. *p-val≤0.05.

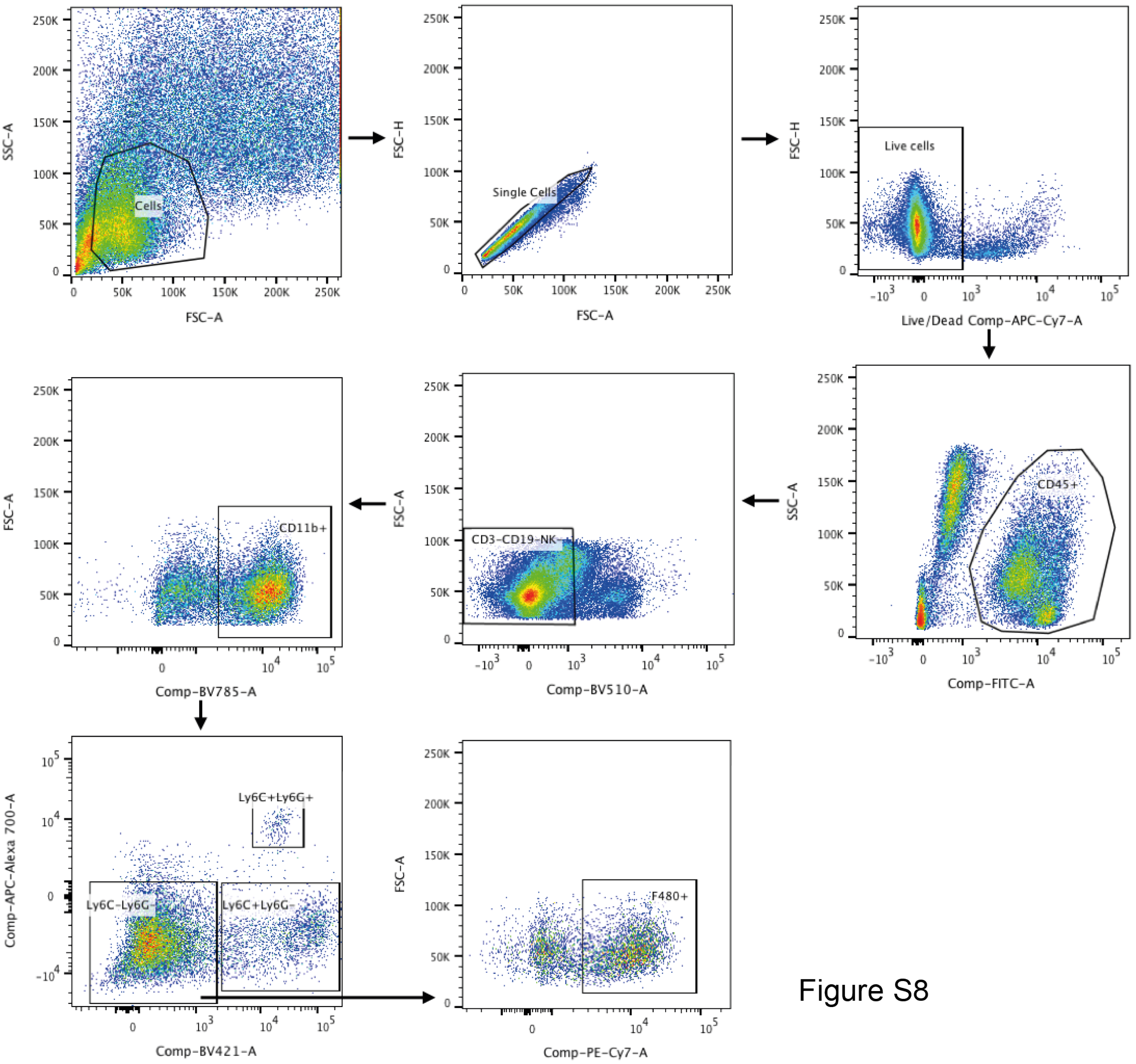


Figure S8

**Supplementary Figure 8: Gating strategy of myeloid infiltrate in subcutaneous tumors and tumor-infiltrating monocytes profile.**

Cells were identified according to FSC and SSC, excluding cell debris. Doublets and cell clumps were then excluded, and live cells were selected. A gate on CD45$^{+}$CD3$^{-}$CD19$^{-}$NK-1.1$^{-}$ allowed the selection of non-lymphoid cells, followed by a gate on CD11b$^{+}$ cells which were divided in Ly6C$^{+}$Ly6G$^{+}$ cells, Ly6C$^{+}$Ly6G$^{-}$ cells and Ly6C$^{-}$Ly6G$^{-}$ cells. Among this last gate were selected F4/80$^{+}$ cells.

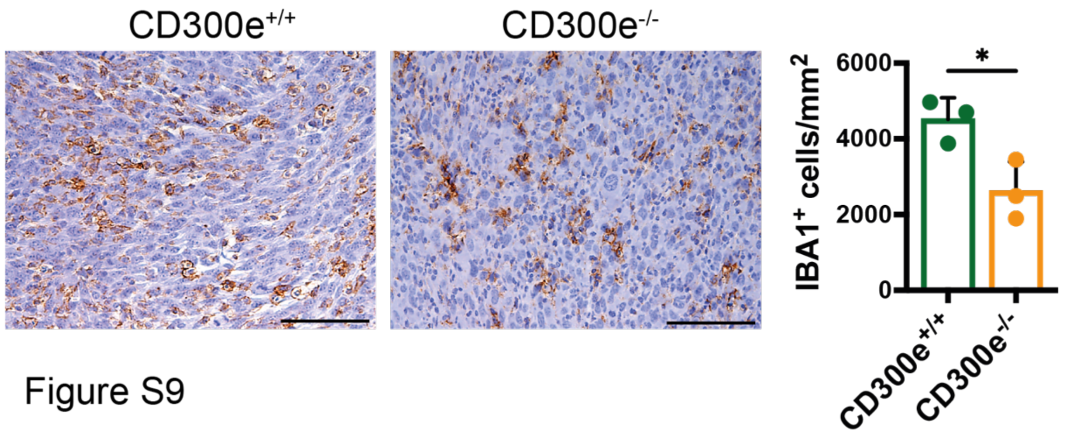


Figure S9

**Supplementary Figure 9: Tumor-infiltrating macrophages in MC38 tumor bearing mice.**

Representative images of paraffin-embedded MC38 tumors of $CD300e^{+/+}$ and $CD300e^{-/-}$ mice immunostained for IBA1 to identify infiltrated macrophages, and relative quantification. Data are expressed as $IBA1^{+}$ cells on $mm^2$, mean ± SEM, n=3 $CD300^{+/+}$ and 3 $CD300e^{-/-}$ mice. Scale bar=100 µm. Statistical significance was determined by unpaired Student's t-test. *p-val≤0.05.

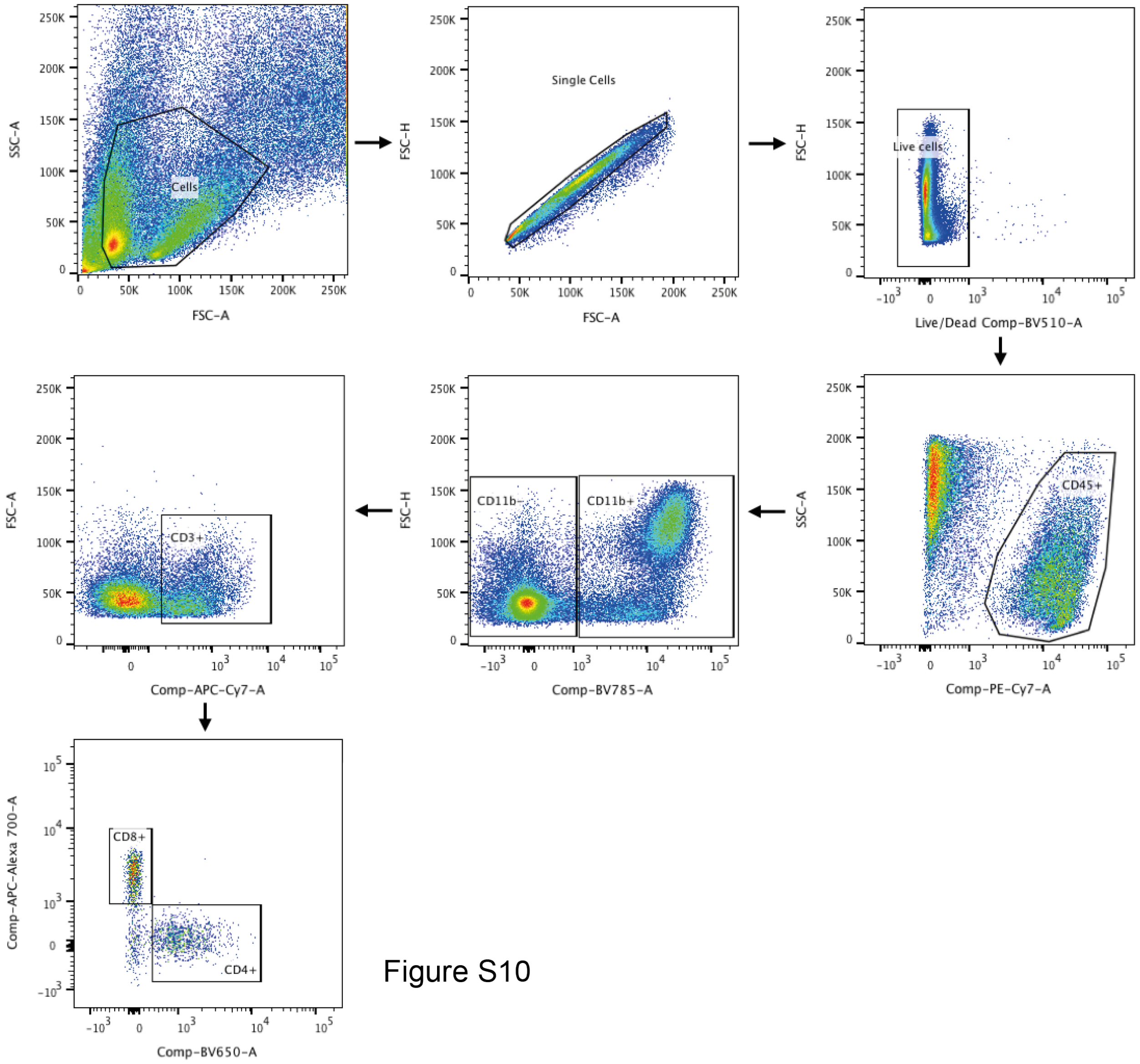


Figure S10

**Supplementary Figure 10: Gating strategy of lymphoid infiltrate in subcutaneous tumors.**

Cells of interest were selected by excluding cell debris, aggregates, and dead cells. $CD3^+$ T lymphocytes were then identified among $CD45^+CD11b^-$ cells. Within $CD3^+$ cells were distinguished between $CD4^+$ and $CD8^+$ cells.

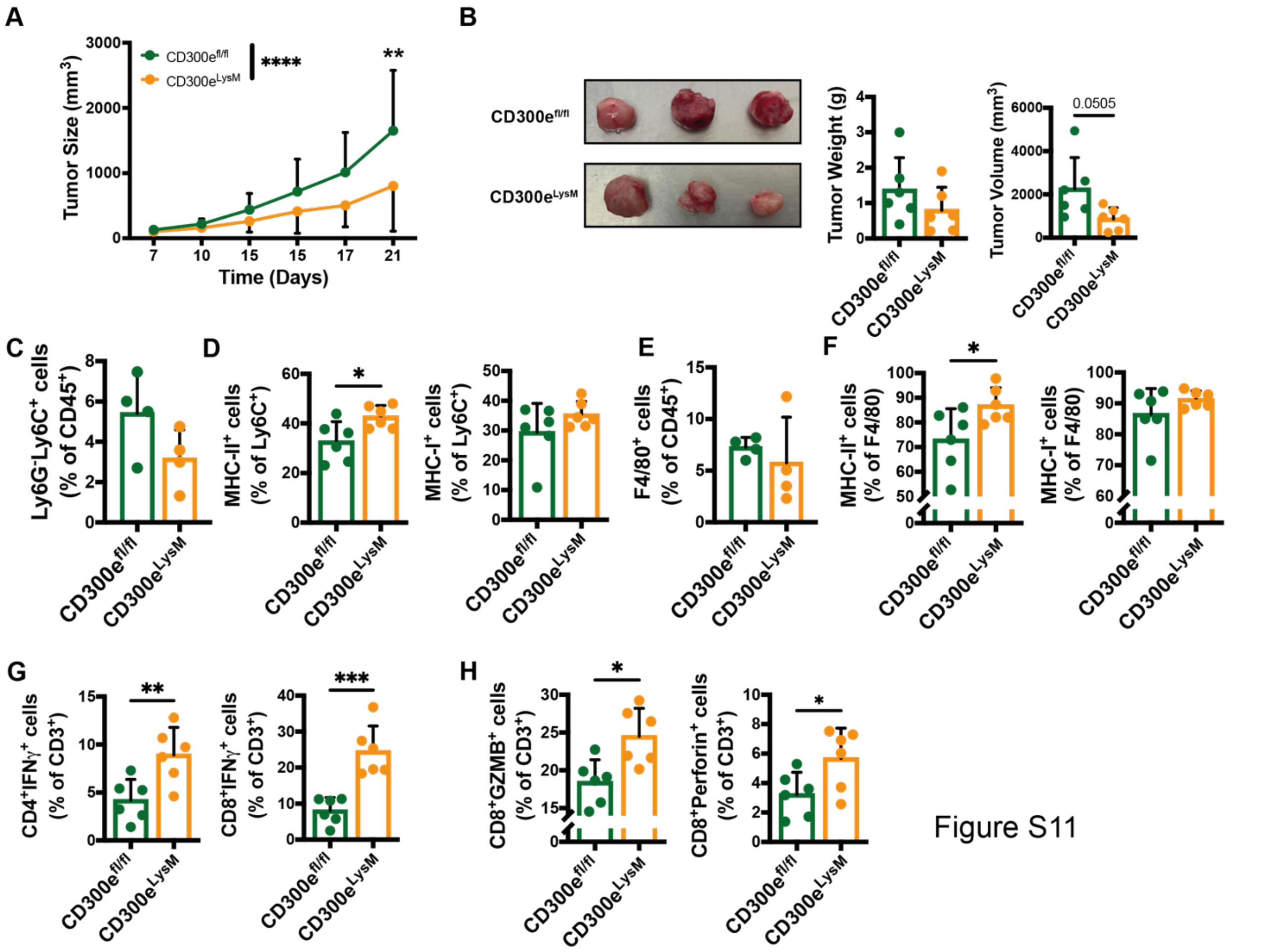


**Supplementary Figure 11: CD300e conditional knock out (CD300e$^{fl/fl}$LysM$^{Cre}$) mice develop a lower tumor burden.**

A) Growth kinetics of MC38 colon adenocarcinoma in conditional KO mice (mean ± SD of n = 6 CD300e$^{fl/fl}$ and 6 CD300e$^{LysM}$ mice). Statistical significance was determined by Two-way ANOVA and Šídák's multiple comparisons test: **p-val≤0.01; **** p-val≤0.0001.

B) Images of tumors (left), tumors' weight (middle) and the final volume (right).

C) Ly6G$^-$Ly6C$^+$ monocytes gated on CD45$^+$CD11b$^+$ cells and shown as percentage of CD45$^+$.

D) MHC-II and MHC-I-expressing tumor-infiltrated monocytes gated on CD45$^+$CD11b$^+$Ly6G$^-$Ly6C$^+$ and shown as percentage of Ly6G$^-$Ly6C$^+$.

E) F4/80$^+$ macrophages gated on CD45$^+$CD11b$^+$Ly6C$^-$ cells and shown as percentage of CD45$^+$.

F) MHC-II and MHC-I-expressing tumor-infiltrated macrophages gated on CD45$^+$CD11b$^+$Ly6C$^-$ F4/80$^+$cells and shown as percentage of F4/80$^+$ cells.

G) IFN-γ production by tumor-infiltrating CD4$^+$ and CD8$^+$ T cells.

H) GZMB and Perforin expression by tumor-infiltrating CD8$^+$ T lymphocytes.

Statistical significance was determined by Student's t test. *p-val≤0.05; **p-val≤0.01; ****p-val≤0.0001.

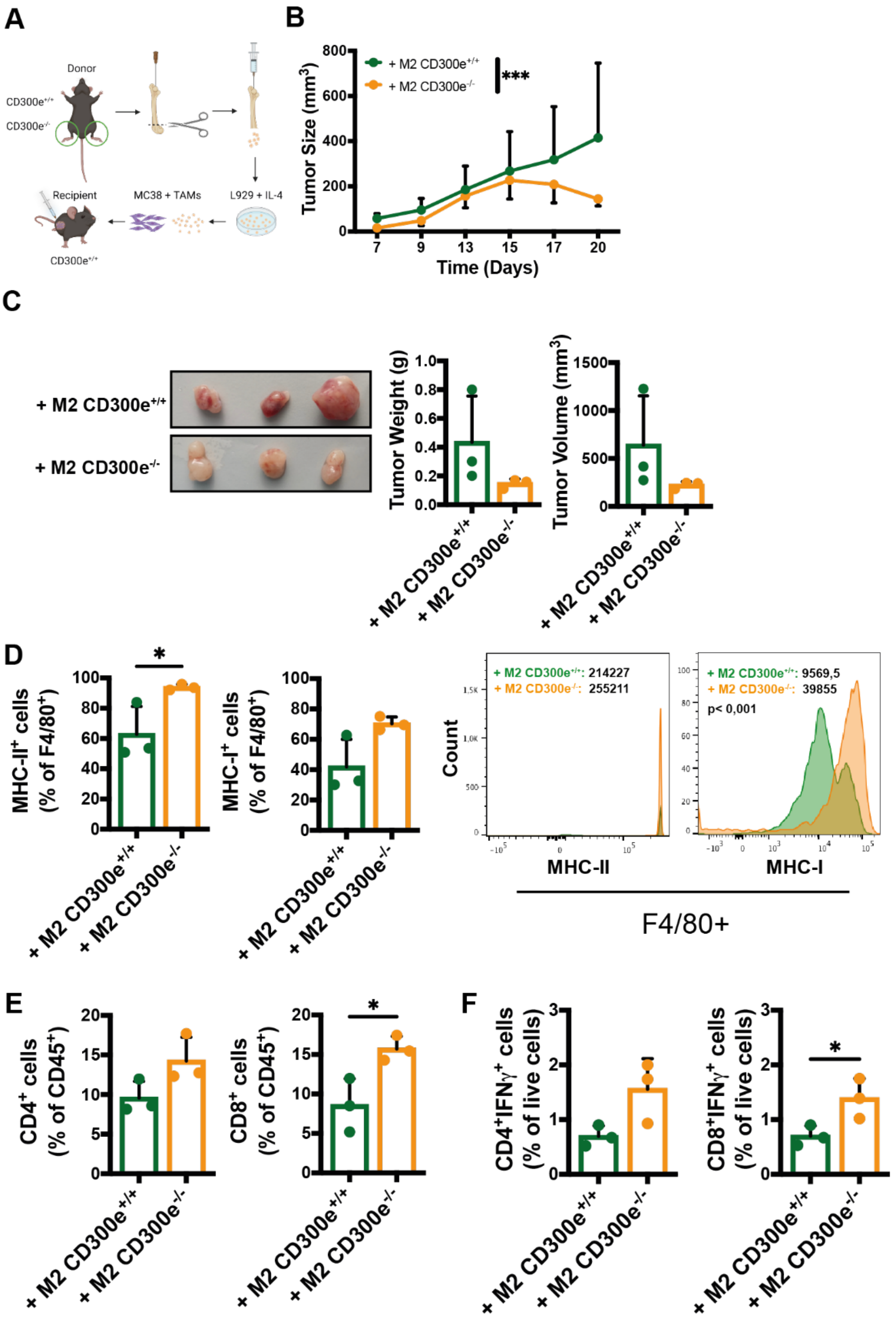
A
Donor
CD300e+/+
CD300e-/-
Recipient
MC38 + TAMs
L929 + IL-4
CD300e+/+
B
+ M2 CD300e+/+
+ M2 CD300e-/-
***
Tumor Size (mm3)
Time (Days)
C
+ M2 CD300e+/+
+ M2 CD300e-/-
Tumor Weight (g)
Tumor Volume (mm3)
D
MHC-II+ cells (% of F4/80+)
MHC-I+ cells (% of F4/80+)
+ M2 CD300e+/+: 214227
+ M2 CD300e-/-: 255211
+ M2 CD300e+/+: 9569,5
+ M2 CD300e-/-: 39855
p< 0,001
Count
MHC-II
MHC-I
F4/80+
E
CD4+ cells (% of CD45+)
CD8+ cells (% of CD45+)
F
CD4+IFNγ+ cells (% of live cells)
CD8+IFNγ+ cells (% of live cells)

Figure S12

**Supplementary Figure S12: BMDM lacking CD300e expression counteract tumor development and impact tumor myeloid and lymphoid landscape.**

BMDMs obtained from CD300e$^{+/+}$ and CD300e$^{-/-}$ mice were stimulated *in vitro* with IL-4 and IL-13 to induce a TAM phenotype, mixed with MC38 cells, and adoptively transferred into CD300e$^{+/+}$ recipient mice.

A) Schematic diagram of macrophage adoptive transfer model based on the s.c. injection of MC38-TAMs-like BMDMs mixture. Created with Biorender.

B) Growth kinetics of MC38 tumors in injected mice (mean ± SD of n = 3 CD300e$^{+/+}$ and 3 CD300e$^{-/-}$ TAMs-transplanted mice). Statistical significance was determined by Two-way ANOVA and Šídák's multiple comparisons test: ***p-val≤0.001

C) Representative pictures of tumors (left), tumor weight (middle) and volume (right) (mean ± SD).

D) MHC-II and MHC-I-expressing tumor-infiltrated macrophages (left) and representative histograms of markers' surface expression (right). Data are expressed as percentage of F4/80$^{+}$ cells ± SD and mean fluorescence intensity (MFI) ± SD, respectively.

E) The frequency of CD4$^{+}$ and CD8$^{+}$ infiltrating T lymphocytes was determined in tumor immune infiltrates from CD300e$^{+/+}$ mice injected with CD300e$^{+/+}$ or CD300e$^{-/-}$ macrophages. Data are expressed as percentage of CD45$^{+}$ cells (mean ± SD).

F) IFN-γ production by tumor-infiltrating CD4$^{+}$ and CD8$^{+}$ T cells (mean ± SD of n = 3 CD300e$^{+/+}$ and 3 CD300e$^{-/-}$ TAMs-transplanted mice).

Statistical significance was determined by Student's t test. *p≤0.05.

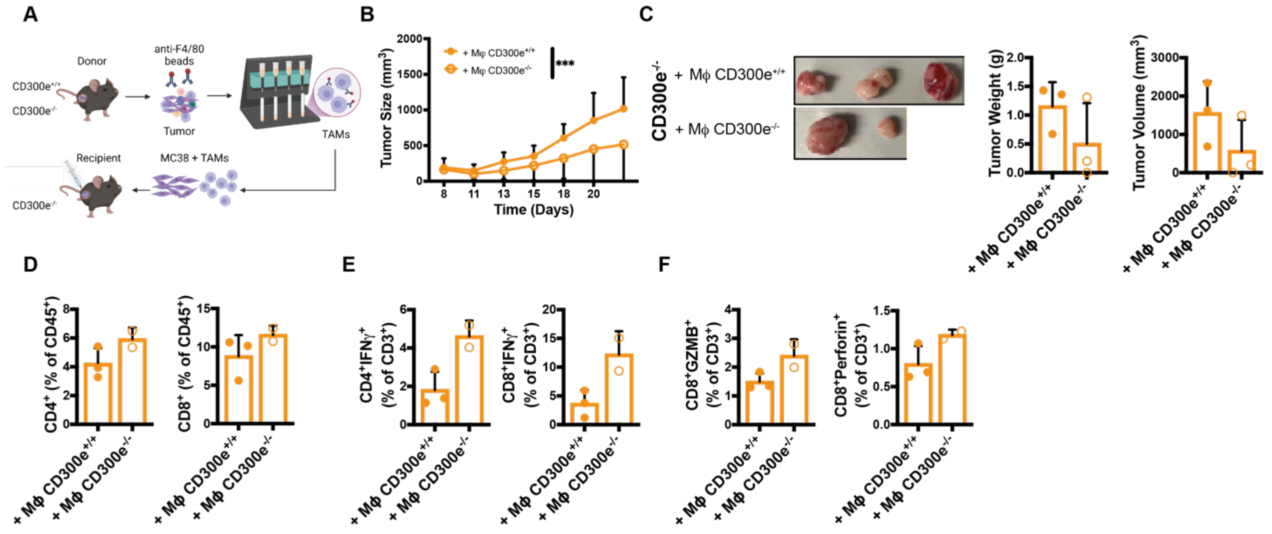


**Supplementary Figure 13: Tumor-infiltrating macrophages lacking CD300e expression are responsible for modulating the tumor microenvironment (CD300e$^{-/-}$ recipient controls).**

A) Schematic representation of experiment workflow. Created with Biorender.

B) Growth kinetics tumor in KO mice (mean ± SD of n = 3 CD300e$^{+/+}$ macrophages and 3 CD300e$^{-/-}$ macrophages). Statistical significance was determined by Two-way ANOVA and Šídák's multiple comparisons test: ***p-val≤0.001.

C) Images of tumors (left), tumors' weight (middle) and the final volume (right).

D) CD4$^{+}$ and CD8$^{+}$ T lymphocytes populations in tumors. Data are expressed as percentage of CD45$^{+}$ cells (mean ± SD).

E) IFN-γ production by tumor-infiltrating CD4$^{+}$ and CD8$^{+}$ T cells Data are expressed as percentage of CD3$^{+}$ cells (mean ± SD).

F) GZMB and Perforin expression by tumor-infiltrating CD8$^{+}$ T lymphocytes. Data are expressed as percentage of CD3$^{+}$ cells (mean ± SD).
